# Comparative biofilm assays using *Enterococcus faecalis* OG1RF identify new determinants of biofilm formation

**DOI:** 10.1101/2021.02.24.432758

**Authors:** Julia L. E. Willett, Jennifer L. Dale, Lucy M. Kwiatkowski, Jennifer L. Powers, Michelle L. Korir, Rhea Kohli, Aaron M. T. Barnes, Gary M. Dunny

## Abstract

*Enterococcus faecalis* is a common commensal organism and a prolific nosocomial pathogen that causes biofilm-associated infections. Numerous *E. faecalis* OG1RF genes required for biofilm formation have been identified, but few studies have compared genetic determinants of biofilm formation and biofilm morphology across multiple conditions. Here, we cultured transposon (Tn) libraries in CDC biofilm reactors in two different media and used Tn sequencing (TnSeq) to identify core and accessory biofilm determinants, including many genes that are poorly characterized or annotated as hypothetical. Multiple secondary assays (96-well plates, submerged Aclar, and MultiRep biofilm reactors) were used to validate phenotypes of new biofilm determinants. We quantified biofilm cells and used fluorescence microscopy to visualize biofilms formed by 6 Tn mutants identified using TnSeq and found that disrupting these genes (OG1RF_10350, *prsA*, *tig*, OG1RF_10576, OG1RF_11288, and OG1RF_11456) leads to significant time- and medium-dependent changes in biofilm architecture. Structural predictions revealed potential roles in cell wall homeostasis for OG1RF_10350 and OG1RF_11288 and signaling for OG1RF_11456. Additionally, we identified growth medium-specific hallmarks of OG1RF biofilm morphology. This study demonstrates how *E. faecalis* biofilm architecture is modulated by growth medium and experimental conditions, and identifies multiple new genetic determinants of biofilm formation.

**Importance:** *E. faecalis* is an opportunistic pathogen and a leading cause of hospital-acquired infections, in part due to its ability to form biofilms. A complete understanding of the genes required for *E. faecalis* biofilm formation as well as specific features of biofilm morphology related to nutrient availability and growth conditions is crucial for understanding how *E. faecalis* biofilm-associated infections develop and resist treatment in patients. We employed a comprehensive approach to analysis of biofilm determinants by combining TnSeq primary screens with secondary phenotypic validation using diverse biofilm assays. This enabled identification of numerous core (important under many conditions) and accessory (important under specific conditions) biofilm determinants in *E. faecalis* OG1RF. We found multiple genes whose disruption results in drastic changes to OG1RF biofilm morphology. These results expand our understanding of the genetic requirements for biofilm formation in *E. faecalis* that affect the time course of biofilm development as well as the response to specific nutritional conditions.

## Introduction

*Enterococcus faecalis* is an early colonizer of the human gastrointestinal (GI) tract, where it remains as a minor component of the healthy microbiota in adults (1–3). It is also a prolific opportunistic pathogen that causes biofilm-associated infections such as infected root canals, bacterial endocarditis, and prosthetic joint infections, and is frequently isolated from polymicrobial infection sites such as the urinary tract, burns, and diabetic foot ulcers (4–9). The ability of *E. faecalis* to thrive as both a commensal and a pathogen is due in part to intrinsic and acquired antibiotic resistance mechanisms, including biofilm formation (10–13). Biofilm development occurs in both the pathogenic and non-pathogenic lifestyles of this organism, and recent high-resolution microscopic analysis of *E. faecalis* biofilms formed in the murine GI tract revealed small matrix-encapsulated microcolonies of biofilm cells spread across the epithelial surface (14). Biofilms formed *in vivo* morphologically resemble those grown *in vitro* (15, 16).

Numerous model systems have been developed to study biofilm formation *in vitro*, including widely used 96-well plate assays, CDC biofilm reactors (CBRs) for assessing biofilms under shear stress and continuous nutrient exchange, and microscopy-based methods that enable fine-scale evaluation of biofilm morphology and matrix properties over a range of time scales (17–19). However, gene expression patterns, biofilm architecture, and genetic determinants of biofilm formation can vary dramatically in biofilms cultured in different model systems, and we have demonstrated that *E. faecalis* biofilm development is influenced by growth medium and nutrient availability (14, 20, 21). Therefore, comparative studies can be useful for understanding how biofilm formation, development, and composition vary across conditions. Incorporation of diverse experimental systems for biofilm growth into the validation of genetic screens using transposon (Tn) libraries may enhance the power of such screens.

Previously, we described the generation of two sequence-defined collections of *E. faecalis* OG1RF Tn mutants termed SmarT (Sequence-defined *mariner* Technology) libraries due to the high level of genetic coverage (insertions in ∼70% of genes and intergenic regions) with a minimal number of Tn mutants (22). SmarT TnSeq library #1 contains 6,829 mutants in genes and intergenic regions. SmarT TnSeq library #2 is a subset of library #1 and contains 1,948 Tn insertions in intergenic regions or uncharacterized or poorly characterized genes (22, 23). These Tn libraries have been used to identify OG1RF genes important for cholic acid resistance, biofilm formation and biofilm-associated antibiotic resistance in microtiter plates, response to phage infection, vaginal colonization, and augmentation of *E. coli* growth (9, 22–27). However, to date no studies have used *E. faecalis* Tn libraries for transposon sequencing (TnSeq) studies to evaluate biofilm fitness determinants comprehensively.

Here, we used a variety of assays for analysis of genetic determinants of OG1RF biofilm formation *in vitro*. Using CBRs, we compared the biofilm fitness of OG1RF Tn mutants in multiple input libraries and in different growth media using TnSeq. We compared these results to previous genetic screens and identified a core set of OG1RF genes required for biofilm formation under multiple conditions. We then measured biofilm formation of a subset of Tn mutants in three secondary biofilm assays (microtiter plates, growth on submerged substrates, and miniature continuous flow biofilm reactors). Additionally, we used bioinformatic tools to predict structure and function for poorly characterized biofilm determinants. Taken together, our data shows that *E. faecalis* OG1RF encodes numerous previously unidentified determinants of biofilm formation, many of which affect biofilm architecture in a temporal and growth medium-dependent manner. Our primary and secondary screening approaches can also guide future studies of biofilm determinants and temporal morphology changes in other organisms.

## Results

### Identification of biofilm determinants in *E. faecalis* using TnSeq

We sought to use the *E. faecalis* OG1RF SmarT libraries (Figure 1A) to evaluate competitive fitness during biofilm formation in CDC biofilm reactors (CBRs) (22). We chose the CBRs for a primary biofilm screen because the system includes continuous flow and medium replacement and a relatively large surface area for biofilm development, decreasing the chance of “bottlenecking” and stochastic loss of mutants. The system also allows for direct, simultaneous comparison of the population distribution of mutants in the planktonic and biofilm states. We used each SmarT library to inoculate CBRs containing either tryptic soy broth without added dextrose (TSB-D) or modified M9 growth medium (MM9-YEG (28)) with ∼10^9^ CFU bacteria. Both media are routinely used to culture *E. faecalis* biofilms (16, 29). Cultures were grown with static incubation (4-6 hr) after which a peristaltic pump was turned at a flow rate of 8 mL/minute (18-20 hr). DNA was isolated from input, planktonic, and biofilm samples, and Tn insertion sites were sequenced in order to determine the relative abundance of Tn mutants (Figure 1B).

**Figure 1.**
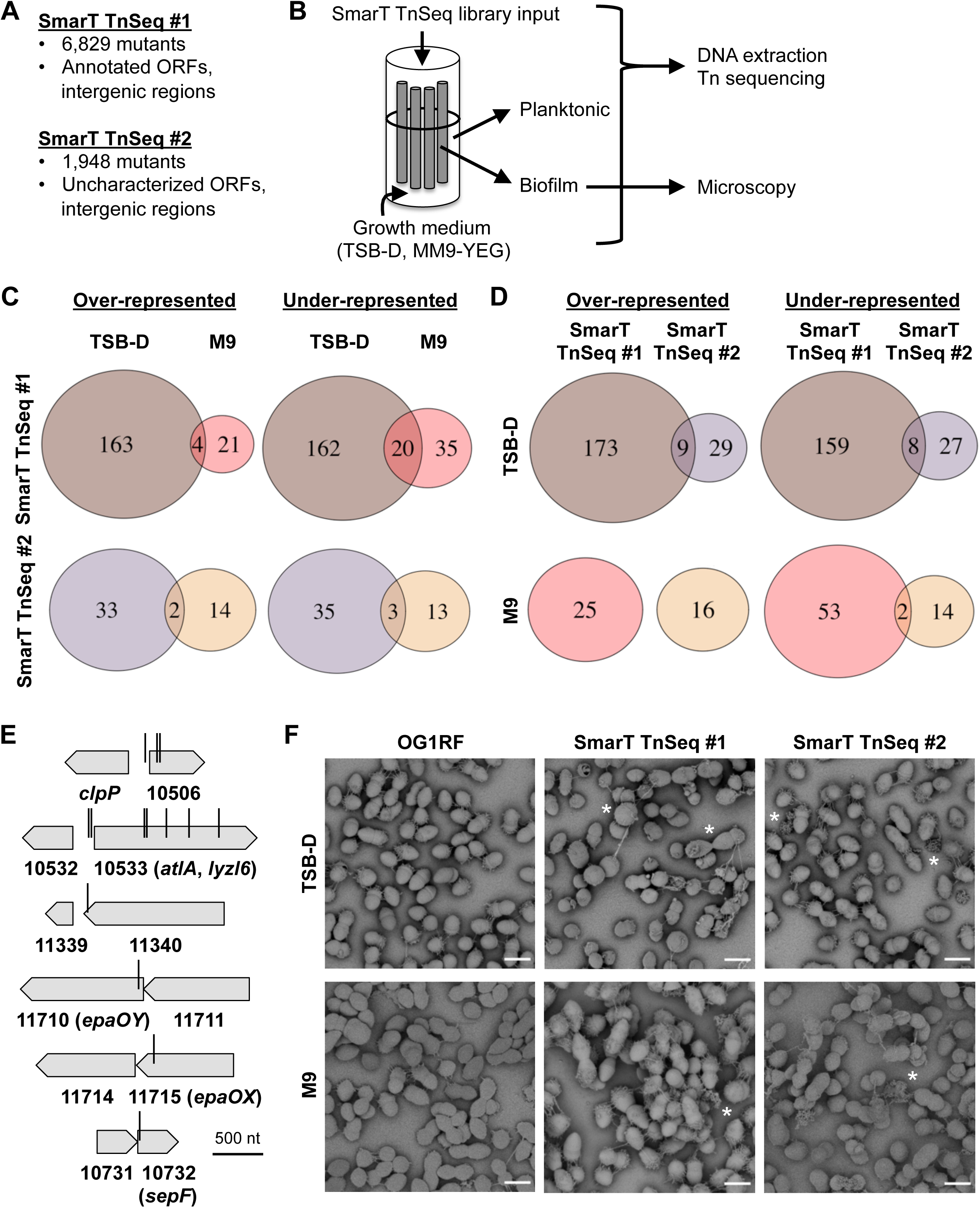
*E. faecalis* OG1RF biofilm formation in CDC reactors and summary of TnSeq. **A)** Summary of SmarT TnSeq libraries used in this study. **B)** Diagram showing CDC biofilm reactor (CBR) inoculation and sampling. **C)** Venn diagrams summarizing differentially abundant (p<0.05, no log_2_FC cutoff) Tn mutants from the same Tn library grown in different media. **D)** Comparison of differentially abundant Tn mutants between both SmarT TnSeq libraries grown in the same media. **E)** Diagrams showing the most underrepresented Tn mutants from biofilm TnSeq. Vertical bars indicate Tn insertion sites. **F)** Scanning electron microscopy images of biofilms from OG1RF and the SmarT TnSeq libraries cultured on Aclar membranes. Examples of misshapen cells and abundant extracellular material are marked with asterisks. Scale bars = 1 μm.

For each medium, we compared Tn abundance between planktonic and biofilm samples to identify mutants over-or underrepresented in biofilms using a significance cutoff of p<0.05 (Figure 1C, Table S1). We first examined Tn mutant abundance in SmarT TnSeq library #1. In TSB-D, 167 mutants were overrepresented and 182 mutants were underrepresented in biofilms relative to planktonic culture (Figure S1A, Figure 1C, brown circles). In MM9-YEG, 25 mutants were overrepresented and 55 mutants were underrepresented in biofilms (Figure S1B, Figure 1C, red circles). Four Tn mutants were overrepresented, and 20 Tn mutants were underrepresented in both TSB-D and MM9-YEG biofilms.

A log_2_ fold change (log_2_FC) of +/- 1.5 was used as a cutoff to identify strongly underrepresented or overrepresented mutants. In TSB-D, 43 mutants had a log_2_FC<-1.5, and 3 had a log_2_FC>1.5. In MM9-YEG, 20 mutants had a log_2_FC<-1.5, and 8 had a log_2_FC>1.5 (Table S1, Figure S1AB). Notably, 13 mutants were strongly underrepresented in both media (Figure 1E, Table 1). These include 2 Tn insertions in OG1RF_10506, a hypothetical gene previously identified in a microtiter plate screen for biofilm-deficient mutants in TSB-D (23), and 5 Tn insertions in *atlA* (OG1RF_10533, *lyzl6*), which encodes a major peptidoglycan hydrolase required for normal cell division and autolysis (30, 31). Additionally, a single Tn insertion in the intergenic region upstream of OG1RF_10506 (named Intergenic_535 based on sequential numbering of intergenic regions in the OG1RF genome) and 2 Tn insertions upstream of *atlA* (Intergenic_563) were underrepresented, suggesting that they could have polar effects on the transcription of OG1RF_10506 and *atlA*. Interestingly, Tn insertions in OG1RF_11710 (*epaOY* (25)) and OG1RF_11715 (*epaOX* (26)) were also strongly underrepresented in biofilms grown in both media. These genes are part of the locus encoding enterococcal polysaccharide antigen (*epa*) (32). Previous work from our laboratory has shown that *epa* genes are associated with biofilm-associated antibiotic resistance, but that Tn insertions in *epa* genes did not lead to reduced biofilm formation in the absence of antibiotics in monoculture (26, 33).

**Table 1.** Tn mutants strongly underrepresented in biofilms grown in both TSB-D and MM9-YEG.

|  |  |  | TSB-D |  | MM9-YEG |  |
| --- | --- | --- | --- | --- | --- | --- |
| Locus tag | Nucleotide position | NCBI description | log<sub>2</sub>FC | P value | log<sub>2</sub>FC | P value |
| Intergenic_535 | 529929 | n/a | -3.23 | 4.18E-13 | -1.56 | 6.01E-27 |
| OG1RF_10506 | 530038 | Hypothetical protein | -2.76 | 2.73E-51 | -1.86 | 1.17E-78 |
| OG1RF_10506 | 530068 | Hypothetical protein | -2.79 | 3.60E-32 | -1.77 | 7.09E-43 |
| Intergenic_563 | 558300 | n/a | -2.69 | 1.53E-3 | -1.62 | 2.11E-4 |
| Intergenic_563 | 558335 | n/a | -3.03 | 7.64E-154 | -1.93 | 8.62E-165 |
| OG1RF_10533 | 559055 | Cell wall lysis protein | -3.19 | 5.89E-159 | -1.58 | 1.26E-102 |
| OG1RF_10533 | 559075 | Cell wall lysis protein | -3.26 | 1.72E-176 | -1.79 | 2.49E-181 |
| OG1RF_10533 | 559358 | Cell wall lysis protein | -3.28 | 8.96E-35 | -1.54 | 3.18E-31 |
| OG1RF_10533 | 559660 | Cell wall lysis protein | -3.19 | 6.76E-108 | -2.22 | 1.44E-122 |
| OG1RF_10533 | 560068 | Cell wall lysis protein | -2.56 | 3.41E-279 | -1.66 | 1.52E-112 |
| OG1RF_11340 | 1403263 | Acetaldehyde dehydrogenase | -2.96 | 1.87E-74 | -1.79 | 2.17E-42 |
| OG1RF_11710 | 1790332 | O-antigen<br>polymerase | -2.12 | 2.90E-4 | -2.36 | 1.28E-14 |
| OG1RF_11715 | 1794475 | Glycosyltransferase | -3.93 | 9.82E-4 | -4.84 | 1.87E-06 |

For SmarT TnSeq library #2, we again used a significance cutoff of p<0.05 to identify Tn mutants differentially represented in biofilms compared to planktonic culture (Table S2, Figure S1). In TSB-D, 35 mutants were overrepresented and 38 mutants were underrepresented in biofilms (Figure S1C, Figure 1C, purple circles). In MM9-YEG, 16 mutants were underrepresented and 16 mutants were overrepresented in biofilms (Figure S1D, Figure 1C, tan circles). Interestingly, we found relatively little overlap when comparing the two libraries in the same medium (Figure 1D). In TSB-D, only 9 of 38 Tn mutants overrepresented in SmarT TnSeq #2 were also overrepresented in SmarT TnSeq #1, and only 8 of 35 Tn mutants underrepresented in SmarT TnSeq #2 were also underrepresented in SmarT TnSeq #1 (Figure 1D, brown and purple circles). There was no overlap of overrepresented mutants in MM9-YEG, and only 2 mutants were underrepresented in both libraries. These results suggest that the community composition affected the relative fitness of Tn mutants in the CBR TnSeq experiments.

Only 4 mutants were underrepresented in SmarT TnSeq library #2 using a log_2_FC cutoff of −1.5, so we used a log_2_FC cutoff of +/-1 to identify strongly under- or overrepresented mutants in this library (Table S2). In TSB-D, 8 mutants had a log_2_FC<-1, including insertions in OG1RF_10506, Intergenic_563, and *bph*, which was previously identified as a phosphatase required for surface attachment and biofilm formation (23). No mutants had a log_2_FC>1. A Tn mutant in OG1RF_10732, which encodes a SepF homolog (34, 35), was strongly underrepresented in both media (Figure 1E). In previous studies, this Tn mutant had varying defects in *in vitro* biofilm formation relative to OG1RF (27, 34), although the specific contribution of SepF to cell division during planktonic and biofilm growth has yet to be reported in *E. faecalis*. The other Tn insertion strongly underrepresented in MM9-YEG is located in Intergenic_1271, which is between OG1RF_11216 and OG1RF_11217. The Tn insertion downstream of Intergenic_1271 in OG1RF_11217 was not underrepresented in either medium, suggesting that Intergenic_1271 may encode a small RNA or peptide that is specifically important for biofilm formation in MM9-YEG.

We also compared biofilms formed by wild type OG1RF versus SmarT TnSeq input pools on Aclar substrates using scanning electron microscopy. Altered biofilm morphology was previously observed in a small pool containing 11 OG1RF Tn mutants in a mouse GI model system (14), and disruption of some *epa* genes led to altered biofilm architecture (16, 33). Parental OG1RF biofilms were visible as a monolayer of cells, with strands of extracellular material present between cells (Figure 1F, left panels). Few cells had aberrant shapes or morphologies. Biofilms formed by the SmarT TnSeq libraries contained markedly more misshapen cells and dysmorphic extracellular material than parental OG1RF biofilms (Figure 1F, center and right panels), suggesting that some Tn insertions in the library disrupt genes involved in cell shape homeostasis or cell division. While additional research is needed to better understand individual determinants of biofilm architecture present in the SmarT TnSeq libraries, these results suggest that both libraries contain a substantial number of mutants with altered cell morphologies that can still form biofilms within complex communities.

### Determination of core and CBR-specific accessory biofilm determinants

In previously reported genetic screens for biofilm determinants, OG1RF Tn mutants were grown as monocultures in microtiter plates (23, 27). This closed, static environment with no competing strains is substantially different than CBRs. To extend our understanding of environmental effects on *E. faecalis* biofilm formation, we sought to determine the overlap between mutants identified from microtiter plate screens and CBR TnSeq, which could constitute core OG1RF biofilm determinants. Because previous screens used TSB-D and not MM9-YEG, we included only the TSB-D TnSeq data sets in this analysis. In previous screens, a total of 204 insertions in 179 genes were associated with statistically reduced biofilm formation (23, 27). Only 35 Tn mutants were identified in both TnSeq and microtiter plate screens (Table 2), including the biofilm-associated phosphatase *bph*, autolysin *atlA*, stress response genes *hrcA* and *dnaK*, and the *ebp* pili operon (23, 36–38).

**Table 2.** Core *E. faecalis* OG1RF biofilm determinants identified in TnSeq and microtiter plate biofilm screens.

| Locus Tag | Nucleotide Position | Gene Name | Description |
| --- | --- | --- | --- |
| Intergenic_442 | 427629 |  | IGR between OG1RF_10412 and OG1RF_10413 |
| Intergenic_464 | 449894 |  | IGR between OG1RF_10434 and OG1RF_10435 |
| OG1RF_10435 | 450277, 450467 | *bph* | Biofilm phosphatase |
| Intergenic_482 | 469369 |  | IGR between OG1RF_10452 and OG1RF_10453 |
| OG1RF_10506 | 530068, 530167, 530274 |  | Hypothetical protein |
| Intergenic_563 | 558335 |  | IGR between OG1RF_10532 and OG1RF_10533 |
| OG1RF_10533 | 559075 | *atlA/lyzI6* | Autolysin, LysM peptidoglycan-binding domain-containing protein |
| OG1RF_10717 | 741838 | *ahrC/argR3* | Arginine repressor |
| OG1RF_10868 | 904848, 905256, 905964 | *ebpR* | M-protein trans acting positive regulator |
| Intergenic_918 | 906315 |  | IGR between OG1RF_10868 and OG1RF_10869 |
| OG1RF_10869 | 906894 | *ebpA* | Endocarditis and biofilm-associated pilus tip protein EbpA |
| OG1RF_10870 | 909926, 910620, | *ebpB* | Endocarditis and biofilm-associated pilus |
|  | 911022 |  | minor subunit EbpB |
| OG1RF_10871 | 911547, 912937 | *ebpC* | Endocarditis and biofilm-associated pilus<br>major subunit EbpC |
| OG1RF_10872 | 913633 | *bps/srtC* | Ebp pilus assembly class C sortase |
| OG1RF_10889 | 928107 | *lepB* | Signal peptidase I |
| Intergenic_1006 | 995480 |  | IGR between OG1RF_10954 and<br>OG1RF_10955 |
| Intergenic_1127 | 1118301 |  | IGR between OG1RF_11075 and<br>OG1RF_11076 |
| OG1RF_11076 | 1118585 | *hrcA* | Heat-inducible transcriptional repressor<br>HrcA |
| OG1RF_11078 | 1120304 | *dnaK* | Molecular chaperone DnaK |
| Intergenic_1130 | 1121988 |  | IGR between OG1RF_11078 and<br>OG1RF_11079 |
| OG1RF_11674 | 1746502 |  | DUF1831 domain-containing protein |
| Intergenic_2022 | 2075283 |  | IGR between OG1RF_11962 and<br>OG1RF_11963 |
| Intergenic_2295 | 2348175 |  | IGR between OG1RF_12228 and<br>OG1RF_12229 |
| OG1RF_12447 | 2581857 |  | DUF3298 domain-containing protein |
| OG1RF_12502 | 2644218 |  | WxL domain-containing protein |
| Intergenic_2613 | 2692363 |  | IGR between OG1RF_r10012 and<br>OG1RF_12535, encodes OG1RF_RS13855 |
| OG1RF_12540 | 2699893 |  | DUF1129 domain-containing protein |

Next, we asked which Tn mutants were underrepresented in biofilm TnSeq but did not have reduced biofilm formation in previous studies. These mutants could have biofilm defects in a community of Tn mutants but not monoculture, or they could be accessory biofilm determinants that are important under flow conditions. Using a log_2_FC cutoff of −1 for the TnSeq results, we identified 55 Tn mutants in 45 genes that were not found in previous studies (Table 3). These include multiple genes in the *epa* operon (OG1RF_11710 (*epaOY*), OG1RF_11714, OG1RF_11715 (*epaOX*), OG1RF_11716, and OG1RF_11722 (*epaQ*)), predicted LCP-family cell wall modifying enzymes (OG1RF_10350, OG1RF_11288), putative transcriptional regulators (OG1RF_12423 and OG1RF_12531), and genes annotated as hypothetical (OG1RF_10968 and OG1RF_11630).

**Table 3.**
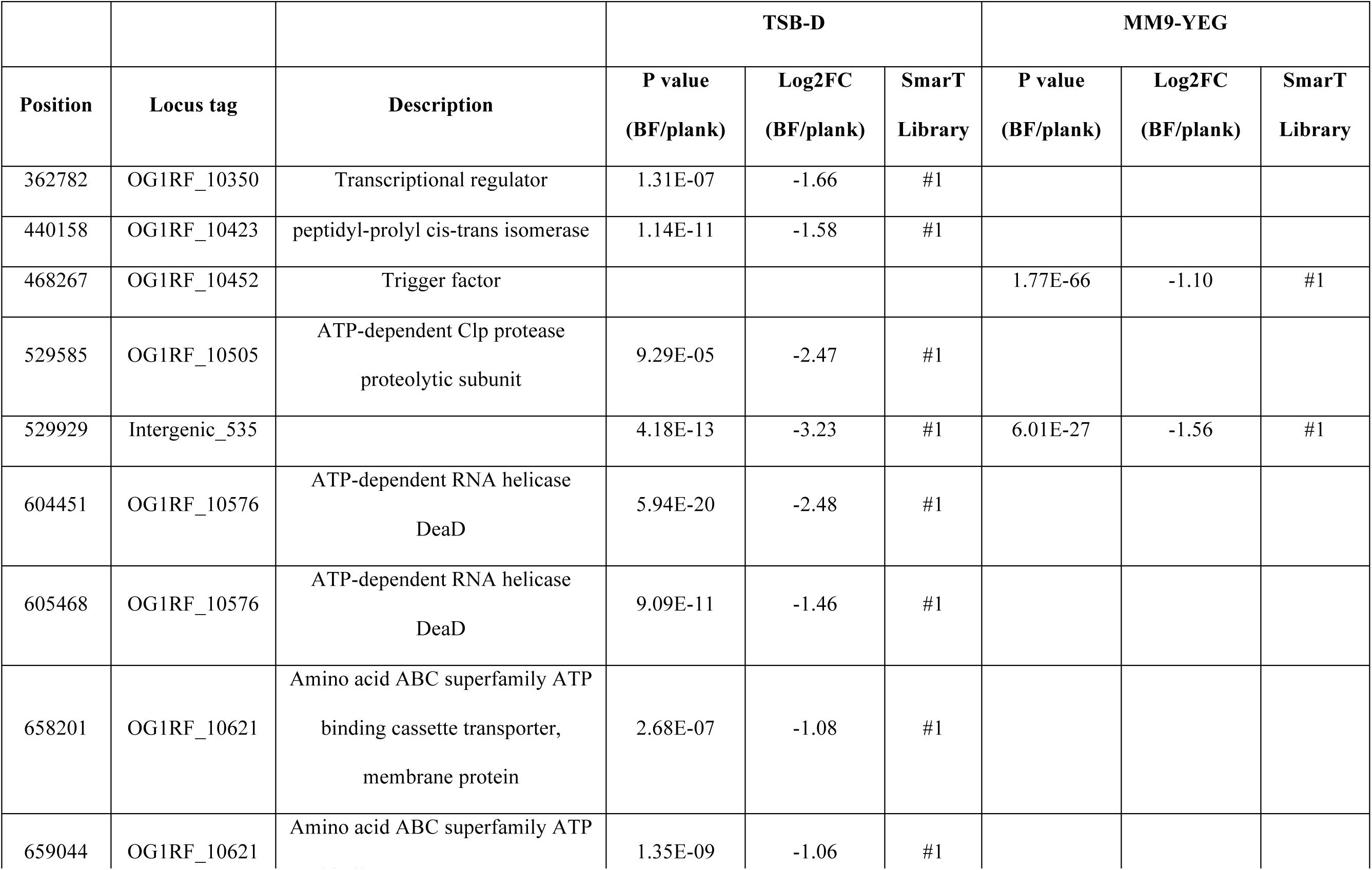
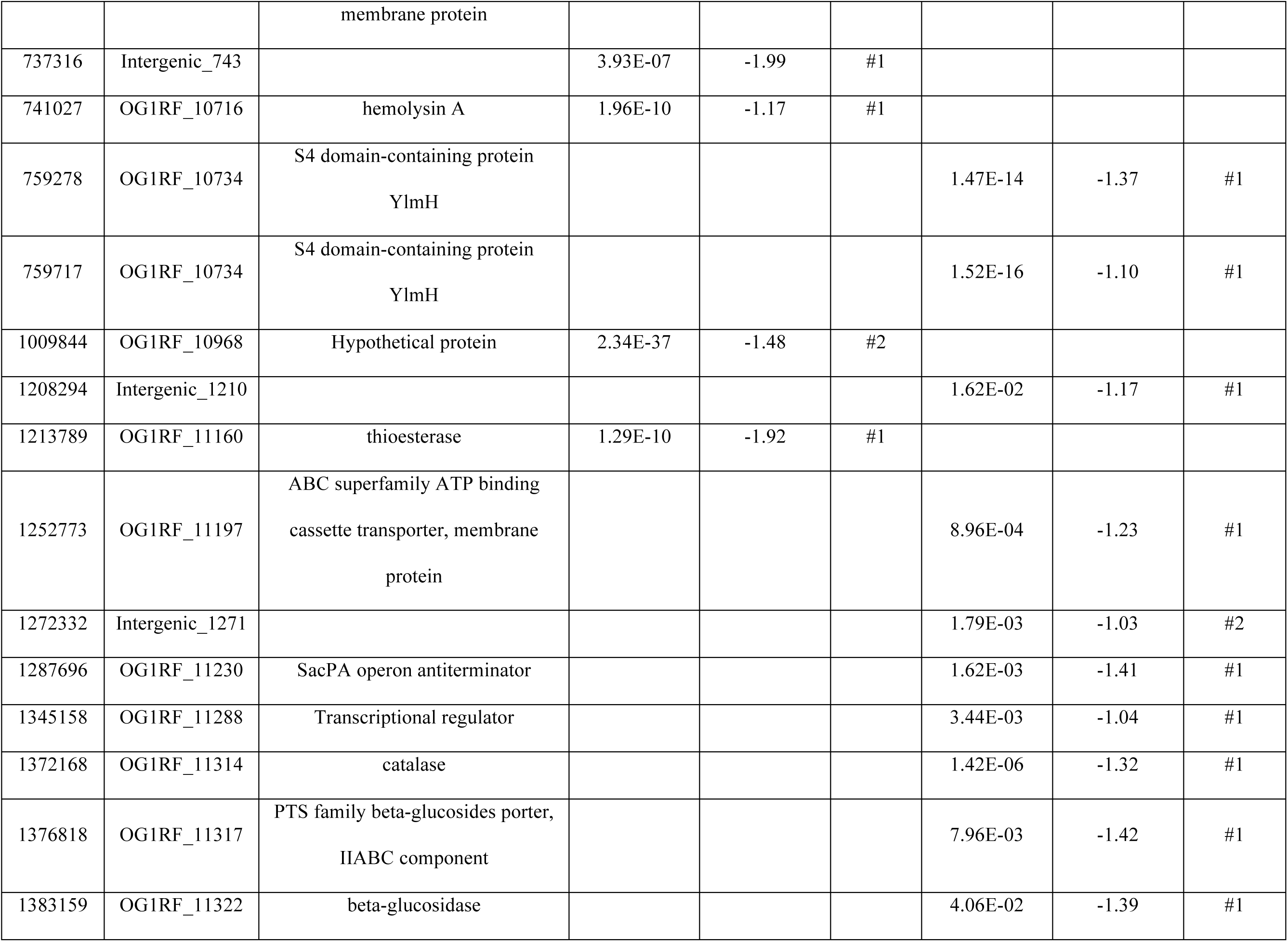
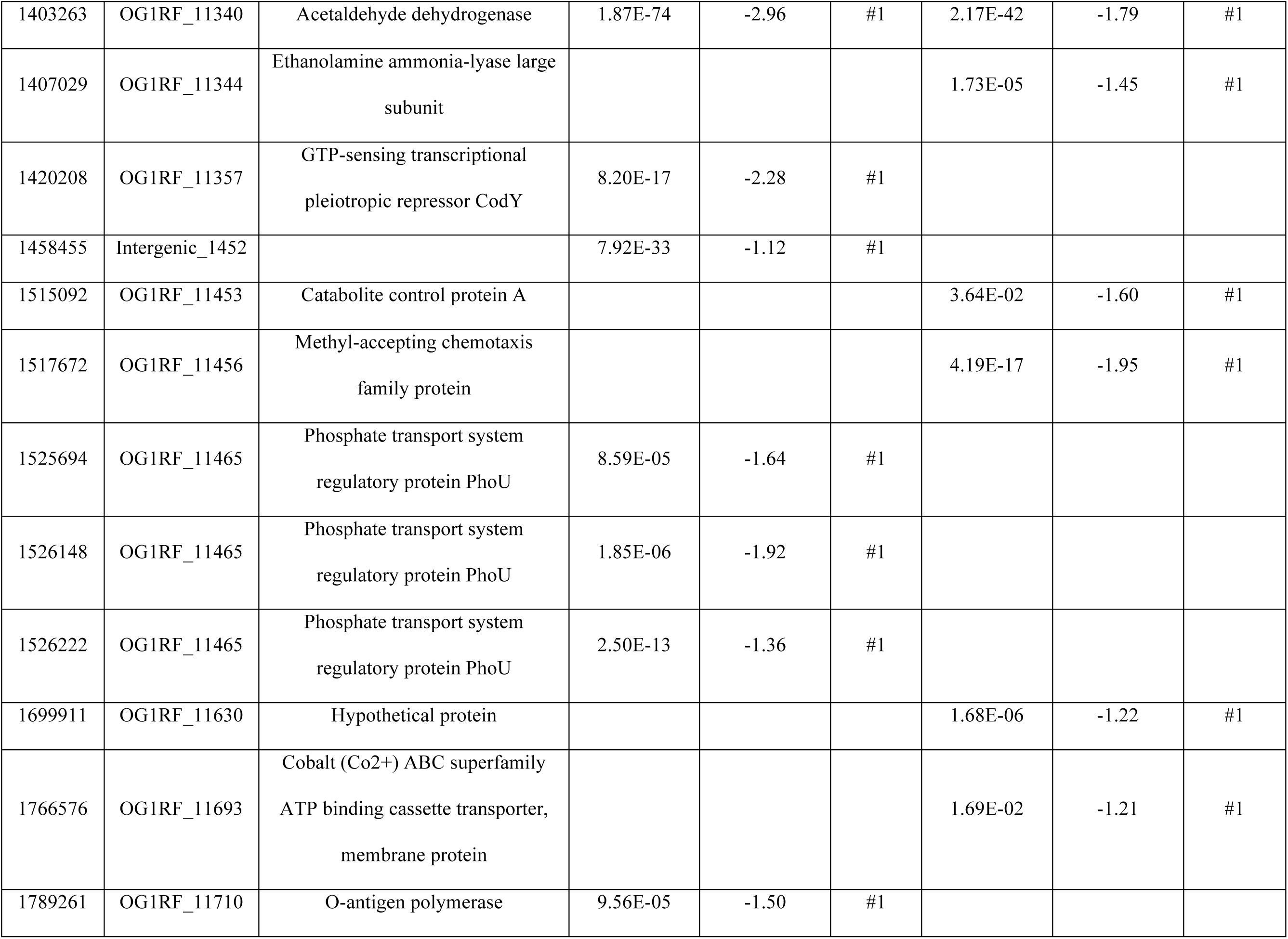
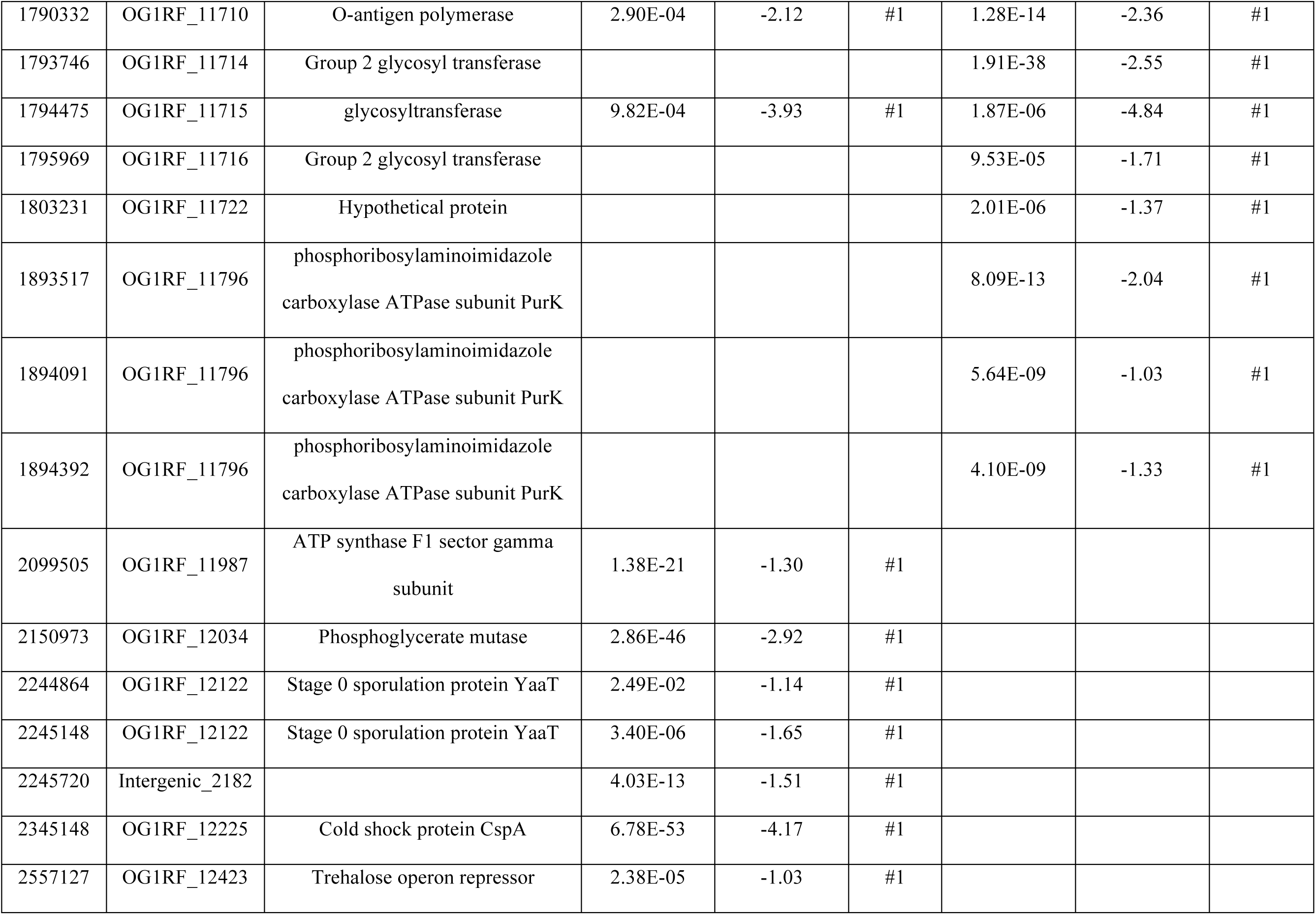
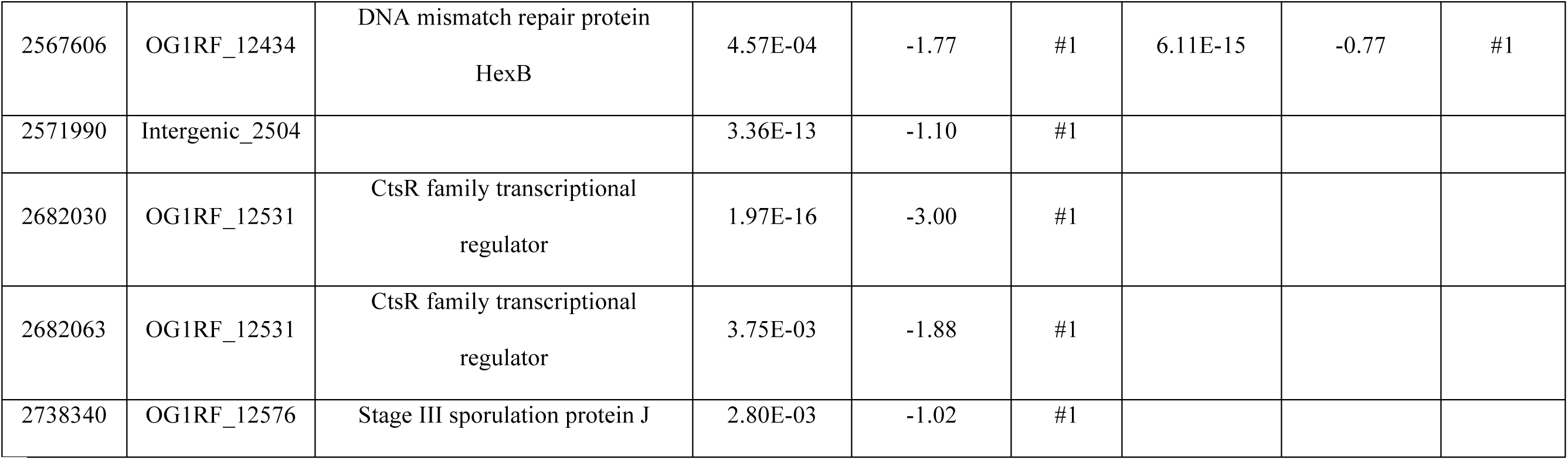
Biofilm determinants not previously identified in genetic screens.

We then sought to validate the importance of these genes for *in vitro* biofilm formation. However, large-scale testing of individual Tn mutants in CBRs is not feasible due to the volume of medium used for each reactor run (∼10 L) as well as the physical size and processing time required for each sample set. Therefore, we chose three previously described *in vitro* experiments to validate biofilm phenotypes: (i) a 96-well plate assay in which biofilm biomass is stained and quantified relative to cell growth (23, 29), (ii) a submerged substrate assay in which biofilms are grown on an Aclar disc covered by growth medium (16, 26), and (iii) a miniature 96-well flow reactor system (MultiRep reactor) in which 96 samples can be cultured in a total of 12 channels on 5 mm disks (33). Because both M9 and TSB-D were used in the CBR TnSeq screen, we carried out the following experiments with both media.

### Phenotypes of “accessory” biofilm determinants in microtiter plate assays

From the 55 Tn mutants presented in Table 3, we obtained 43 Tn mutants from the arrayed SmarT library stock plates. When multiple Tn insertions in a gene were identified, we chose only the insertion closest to the start codon. Additional mutants were excluded based on their location upstream of known biofilm determinants and the possibility that these insertions had polar effects on previously studied genes. To maintain consistency with previous experiments, we measured biofilm production of the Tn mutants at 6 hr and 24 hr. A strain lacking *bph*, previously implicated in biofilm development (23), was used as a negative control, and biofilm production was normalized to OG1RF (Figure 2A). In TSB-D, 12 mutants had significantly altered biofilm production relative to OG1RF at 6 hr (12 decreased, 0 increased) (Figure 2B, black bars), and 5 mutants had altered biofilm levels at 24 hr (3 decreased, 2 increased) (Figure 2B, pink bars). In MM9-YEG, 7 Tn mutants had altered biofilm production at 6 hr (2 decreased, 5 increased) (Figure 2C, black bars), and 6 mutants had altered biofilm levels at 24 hr (5 decreased, 1 increased) (Figure 2C, pink bars). Overall, ∼30% of mutants (13/43) had reduced biofilm formation relative to OG1RF. Interestingly, some mutant strains had higher biofilm production in MM9-YEG than TSB-D, including Δ*bph* and OG1RF_10576, demonstrating that growth medium influences which genes are required for biofilm formation. We did not observe a correlation between the change in abundance (log_2_FC) of Tn mutants in TnSeq and biofilm index in microtiter plate biofilm assays (Figure S1E-H).

**Figure 2.**
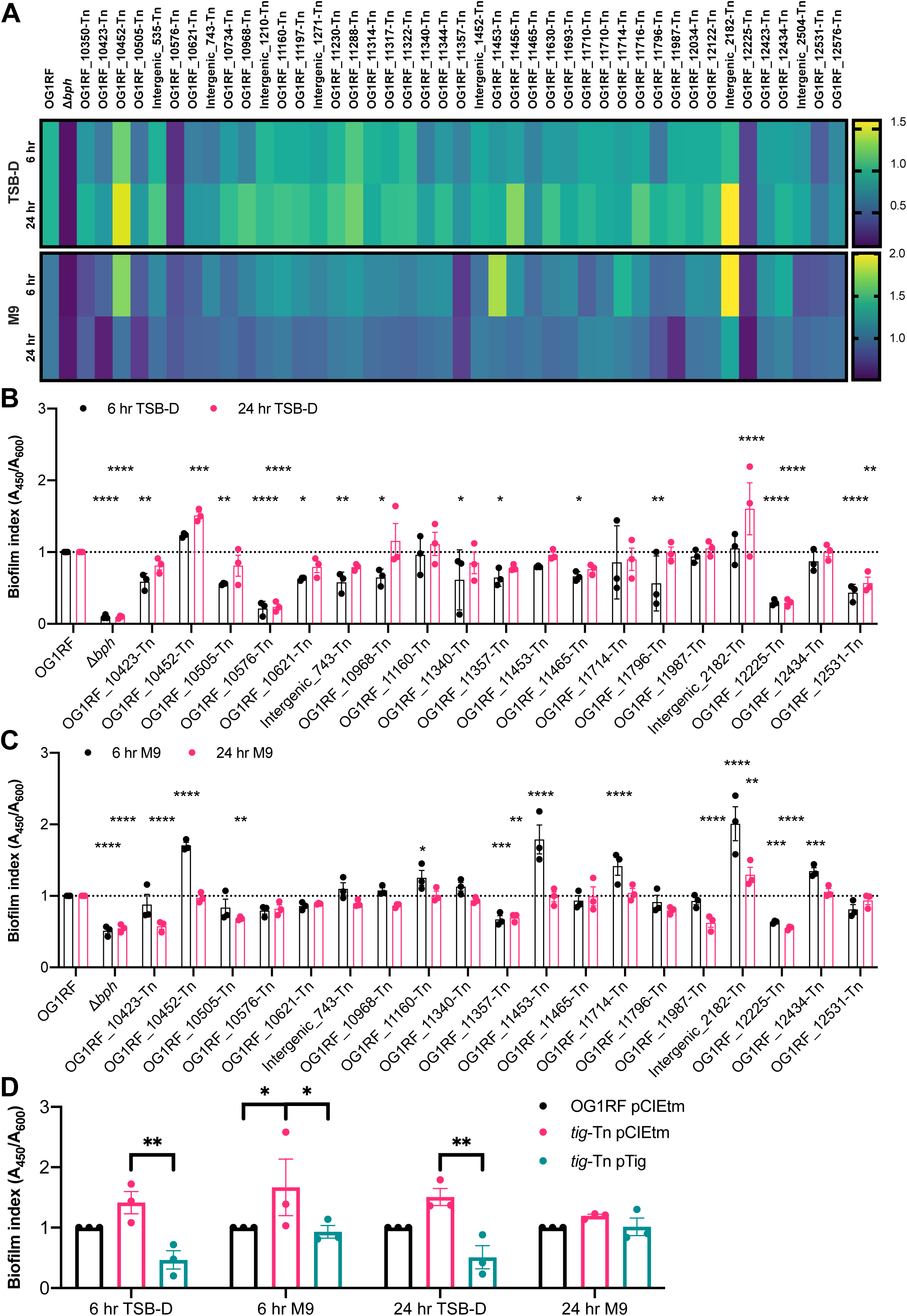
Tn mutants identified from biofilm TnSeq have variable biofilm production in microtiter plates. **A)** Heatmap summarizing biofilm index values (A_450_/A_600_ relative to OG1RF) for all mutants. Biofilm index shading legends are shown on the right. **B)** TSB-D biofilm index values and **C)** MM9-YEG biofilm index values for all Tn mutants with significantly altered biofilm production in either media. For clarity, a dotted line is shown at the OG1RF biofilm index value. Plotted values are the same ones represented in the heat maps in panel **A**. **D)** Biofilm phenotypes were complemented for *tig*-Tn. Strains carried either an empty pCIEtm plasmid or pCIEtm with the wild-type allele cloned under a pheromone-inducible promoter. Biofilm assays were carried out in the growth medium and for the length of time indicated in x-axis labels. All cultures were grown with 25 ng/mL cCF10 to induce expression of the cloned *tig* gene. For panels **BC**, three biological replicates were performed, each with two technical replicates. Statistical significance was evaluated by two-way ANOVA with Dunnett’s multiple comparisons test (*p<0.05, **p<0.01, ***p<0.001, ****p<0.0001). For panels **DE**, three biological replicates were performed, each with three technical replicates. Statistical significance was evaluated by two-way ANOVA with Sidak’s multiple comparisons test (*p<0.05, **p<0.01, ***p<0.001, ****p<0.0001).

Although all 43 Tn mutants were underrepresented in biofilm TnSeq, ∼14% (6/43) had increased biofilm levels relative to OG1RF in 96-well plates (Figure 2BC). We chose to complement the high biofilm phenotype of *tig*-Tn (OG1RF_10452-Tn) by expression of the wild-type gene from a pheromone-inducible plasmid (23). *tig* encodes trigger factor, a chaperone involved in folding newly synthesized proteins (39). Expression of *tig* from a plasmid significantly decreased biofilm relative to the Tn mutant carrying an empty vector plasmid (Figure 2D). The opposing biofilm phenotypes observed for some Tn mutants in CBR TnSeq compared to 96-well plates underscores how determinants of biofilm formation may vary across experimental platforms and suggests that molecular changes during biofilm development are highly sensitive to specific assay conditions.

### Biofilm formation of Tn mutants in submerged substrate assays

We chose 6 of the 43 Tn mutants described above for biofilm assays using submerged Aclar assays, in which strains are cultured in multi-well plates containing Aclar coupons. These permit sampling of both planktonic and biofilm cells for visualization via microscopy and CFU quantification (16, 29). All 6 mutants were underrepresented in at least one library in biofilm TnSeq (Table 3, Table S1, Table S2) but had a range of phenotypes in the microtiter plate assays described above. Relative to parental OG1RF biofilm levels in 96-well plates, *prsA*-Tn (encoding an extracellular peptidyl-prolyl isomerase) and OG1RF_10576-Tn (encoding a predicted DEAD-box helicase) had decreased biofilm. *tig*-Tn (encoding trigger factor) had increased biofilm, and OG1RF_10350-Tn, OG1RF_11456-Tn, and OG1RF_11288-Tn did not have significantly different levels of biofilm compared to OG1RF (Figure 2BC).

We inoculated strains at 10^7^ CFU/mL and quantified planktonic and biofilm CFU after 6 hr. In TSB-D, *prsA*-Tn, OG1RF_10576-Tn, and OG1RF_11456-Tn had significantly lower planktonic CFU/mL than OG1RF (Figure 3A, pink bars). OG1RF_10576-Tn had a ∼1 log decrease in biofilm CFU relative to OG1RF (Figure 3A, green bars), although this difference was not statistically significant. To determine whether mutants had a biofilm-specific decrease in viable cells (as opposed to lower biofilm growth due to growth defects in planktonic culture), we calculated the ratio of biofilm growth to planktonic growth relative to OG1RF. By this metric, only the Δ*bph* strain had a significant reduction relative to OG1RF (Figure 3B).

**Figure 3.**
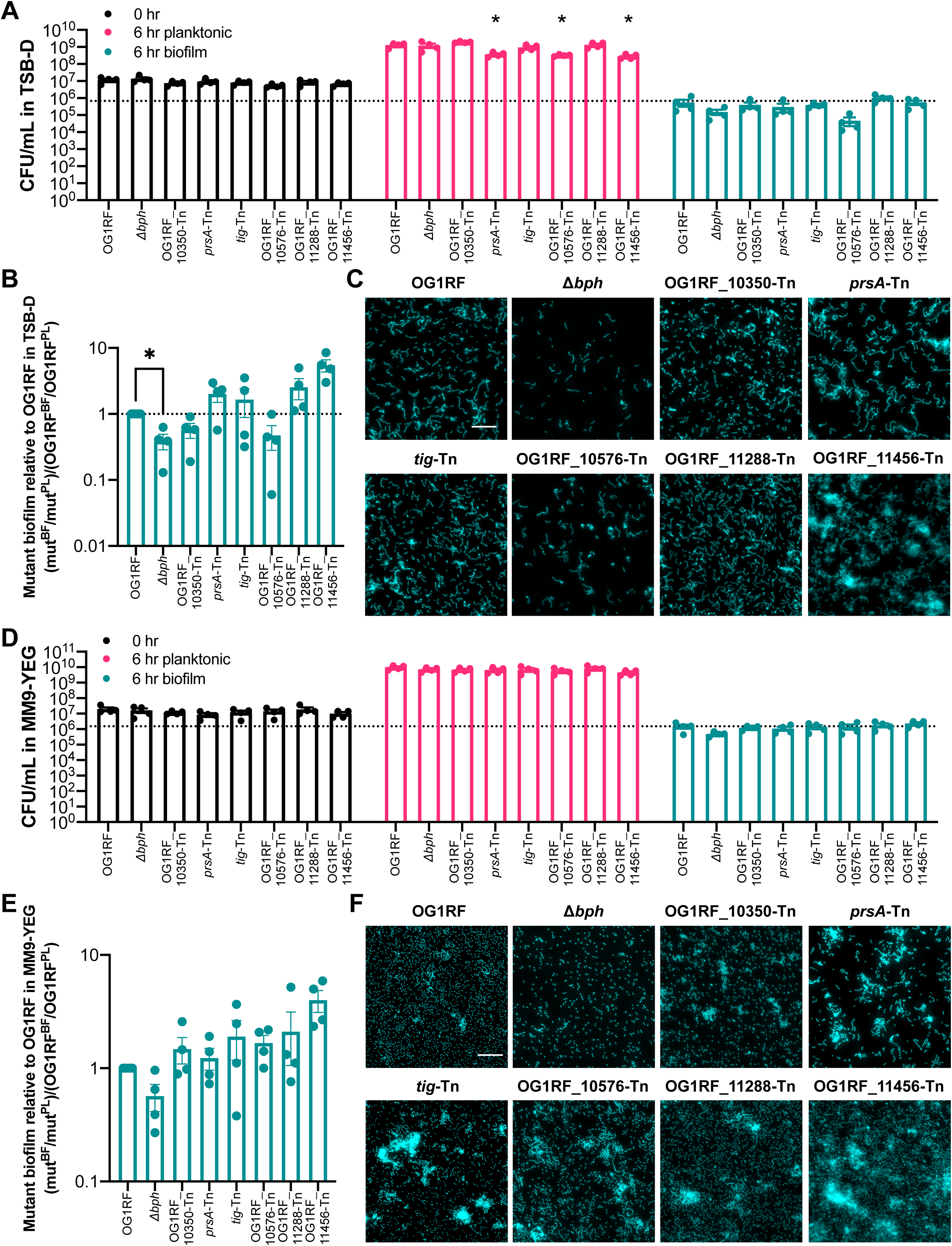
Biofilm formation of selected Tn mutants using submerged Aclar assay. **A)** CFU of strains at 0 hr and 6 hr in TSB-D. The dotted line indicates OG1RF biofilm CFU. **B)** Ratio of biofilm to planktonic growth relative to OG1RF. **C)** Representative microscopy images of Hoechst 33342-stained biofilms from TSB-D cultures. **D)** CFU of strains at 0 hr and 6 hr in MM9-YEG. The dotted line indicates OG1RF biofilm CFU. **E)** Ratio of biofilm to planktonic growth relative to OG1RF. **F)** Representative microscopy images of Hoechst 33342-stained biofilms from MM9-YEG cultures. For panels **A** and **D**, each data point represents the average of two technical replicates, and a total of four biological replicates were performed. Statistical significance was evaluated by two-way ANOVA with Dunnett’s multiple comparisons test (*p<0.05, **p<0.01, ***p<0.001, ****p<0.0001). For panels **B** and **E**, values were obtained using the data points presented in panels A and D, respectively. Statistical significance was evaluated by one-way ANOVA with Dunnett’s multiple comparisons test (*p<0.05, **p<0.01, ***p<0.001, ****p<0.0001). For panels **C** and **F**, samples were grown in parallel to cultures used to generate panels **A** and **D**. Scale bars = 20 μm. Two technical replicates were processed for each biological replicates, and representative images are shown.

Biofilms were visualized with fluorescence microscopy after staining with Hoechst 33342, a nucleic acid label. OG1RF biofilms consistently grew as a monolayer of short chains of bacteria with few multi-cellular aggregates or clumps (Figure 3C). As previously observed, biofilms formed by the Δ*bph* negative control strain contained fewer cells than OG1RF (23). The appearance of OG1RF_10350-Tn, *tig*-Tn, and OG1RF_11288-Tn biofilms was similar to OG1RF. Although there was not a significant reduction in OG1RF_10576-Tn biofilm CFU relative to OG1RF (Figure 3B), these mutant biofilms had visibly less surface coverage than OG1RF biofilms. *prsA*-Tn biofilms contained some multicellular aggregates, and OG1RF_11456-Tn biofilms had large clumps of cells (Figure 3C).

We next examined the growth of these mutants in MM9-YEG. Unlike the corresponding experiments in TSB-D (Figure 3A, pink bars), no mutants had reduced CFU in planktonic culture (Figure 3D, pink bars). Additionally, none of the mutants had reduced CFU in biofilms (Figure 3D, green bars) or the ratio of biofilm to planktonic growth relative to OG1RF (Figure 3E). However, visualization of Aclar substrates revealed substantial differences in biofilm architecture. In MM9-YEG, OG1RF formed a monolayer biofilm composed mainly of single cells and some small aggregates (Figure 3F). The Δ*bph* biofilm had less surface coverage but was still composed of mostly single cells. All Tn mutants formed biofilms with multicellular aggregates. *prsA*-Tn, *tig*-Tn, and OG1RF_10576-Tn biofilms had mixtures of single cells and small multicellular chains, while nearly all cells in OG1RF_11456-Tn biofilms grew as chains and aggregates. Interestingly, fewer multi-cellular chains and more individual cells were observed in biofilms grown in MM9-YEG compared to TSB-D (compare Figure 3C to Figure 3F). Conversely, more large multicellular aggregates were observed in MM9-YEG compared to TSB-D, suggesting that nutritional components could regulate cell chaining and aggregate formation as separate processes during biofilm growth.

### Biofilm formation in miniature flow reactors

MultiRep reactors are miniaturized 12-channel biofilm flow reactors that permit simultaneous sampling of planktonic cultures and biofilms formed on removable Aclar coupons that rest in wells in each channel (Figure S2A). OG1RF biofilms from MultiRep reactors resemble the monolayer biofilms formed in CBRs (14, 16, 33). The same 6 Tn mutant cultures used for submerged Aclar assays in the previous section were inoculated into the MultiRep reactors at 10^7^ CFU/mL and grown with static incubation for 4 hr, after which medium flowed through each channel at a rate of 0.1 mL/min for 20 hr (t_total_ = 24 hrs). The flow rate for growth medium was chosen for consistency in turnover rate compared to CBR experiments. Planktonic and biofilm cultures were quantified and visualized at 4 hr and 24 hr. After 4 hr growth in TSB-D, planktonic cultures of *tig*-Tn, OG1RF_10576-Tn, and OG1RF_11456-Tn had significantly reduced CFU/mL relative to OG1RF (Figure 4A, pink bars). The Δ*bph* negative control strain had significantly reduced biofilm CFU relative to OG1RF, as did *prsA*-Tn, OG1RF_10576-Tn, and OG1RF_11456-Tn (Figure 4A, green bars). However, only Δ*bph* had a biofilm-specific reduction in growth relative to OG1RF at 4 hr (Figure 4B). *tig*-Tn, which had increased biofilm formation in microtiter plate assays (Figure 2D), had a biofilm-specific 1.89-fold increase CFU relative to OG1RF (Figure 4B).

**Figure 4.**
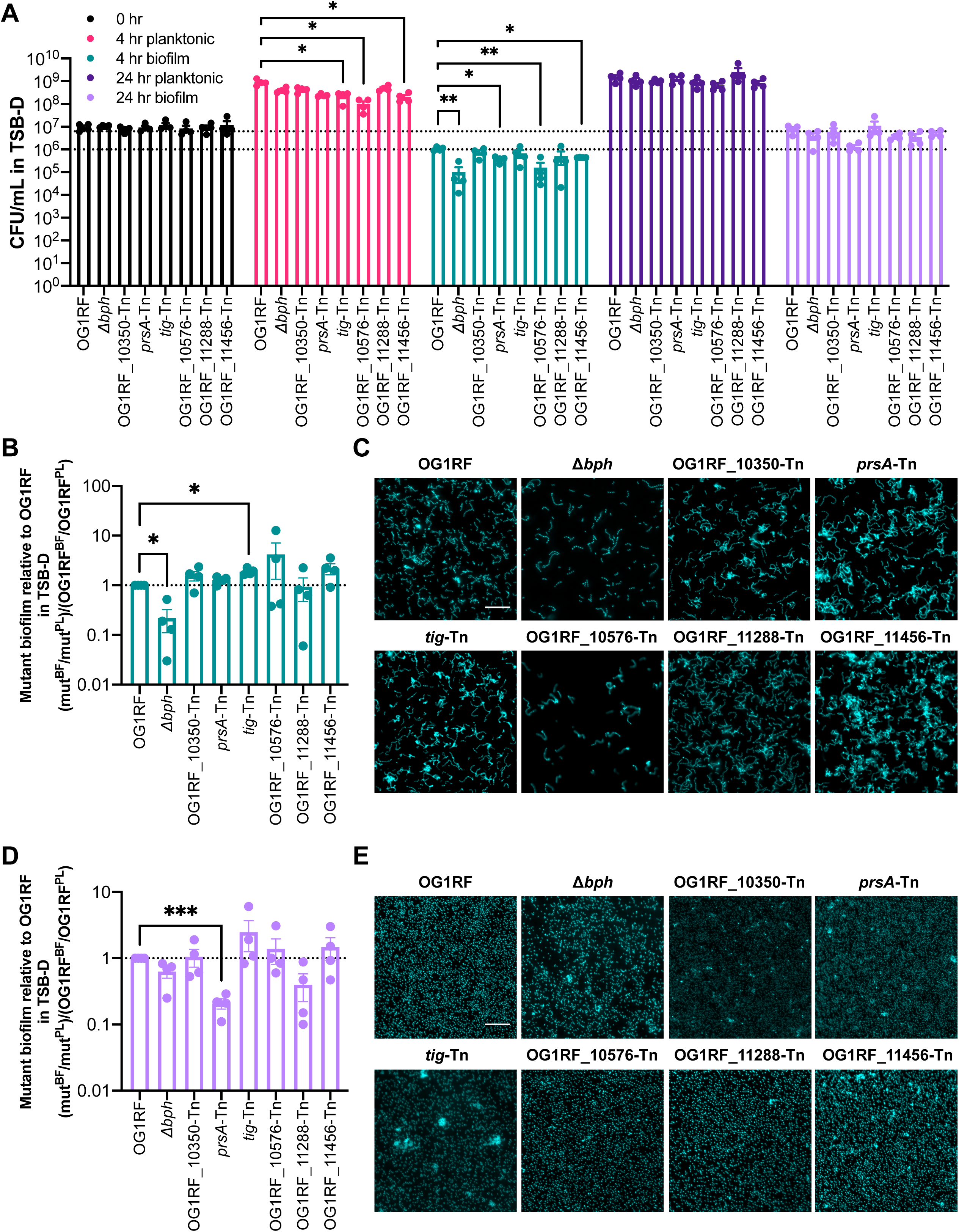
Biofilm formation of selected Tn mutants grown in MultiRep reactors in TSB-D. **A)** CFU of strains at 0 hr, 4 hr, and 24 hr. The dotted lines indicated OG1RF biofilm CFU at 24 hr (top line) and 4 hr (bottom line). **B)** Ratio of biofilm to planktonic growth at 4 hr relative to OG1RF. **C)** Representative microscopy images of Hoechst 33342-stained biofilms at 4 hr. **D)** Ratio of biofilm to planktonic growth at 24 hr relative to OG1RF. **E)** Representative microscopy images of Hoechst 33342-stained biofilms at 24 hr. For panel **A**, each data point represents the average of two technical replicates, and a total of four biological replicates were performed. For panels **B** and **D**, data points were derived using the data points shown in panel **A**. Statistical significance was evaluated by one-way ANOVA with Dunnett’s multiple comparisons test (*p<0.05, **p<0.01, ***p<0.001, ****p<0.0001). For panels **C** and **E**, samples were grown in parallel to cultures used to generate panel **A**. Scale bars = 20 μm. Two technical replicates were processed for each biological replicates, and representative images are shown.

Biofilm appearance was evaluated using fluorescence microscopy of Hoescht 33342-stained cells. After 4 hr, OG1RF formed biofilms with single cells and multi-cell chains but few large aggregates (Figure 4C). Biofilms formed by Δ*bph* and OG1RF_10576-Tn had very few cells, in agreement with the average reduction in biofilm CFU at 4 hr. OG1RF_10350-Tn and *tig*-Tn formed biofilms with chained cells and small clumps, and *prsA*-Tn and OG1RF_11456-Tn formed biofilms with larger clumps of cells. OG1RF_11288-Tn formed biofilms that resembled OG1RF.

After 24 hr, no mutants had reduced planktonic CFU/mL relative to OG1RF (Figure 4A, dark purple bars). Although the biofilm CFU of *prsA*-Tn was ∼1 log lower than OG1RF (Figure 4A, lilac bars), this difference was not statistically significant. However, *prsA*-Tn had a significant reduction in the ratio of biofilm to planktonic cells relative to OG1RF (Figure 4D). In contrast to biofilm morphology at 4 hr, OG1RF biofilms at 24 hr appeared as smooth layers of single cells, and chaining and clumping were not evident (Figure 4E, Figure S2B). Unlike 4 hr biofilms formed by Δ*bph* and OG1RF_10576-Tn, biofilms after 24 hr growth covered most of the Aclar surface. OG1RF_10350-Tn and OG1RF_11288-Tn biofilms resembled OG1RF, and small clumps of cells were visible in *prsA*-Tn, *tig*-Tn, and OG1RF_11456-Tn biofilms.

In MM9-YEG, no mutants had statistically different planktonic or biofilm CFU compared to OG1RF after 4 hr static growth (Figure 5A, pink and green bars) or an additional 20 hr growth under flow conditions (Figure 5A, purple and lilac bars). We observed more variability in planktonic growth of each Tn mutant after 24 hr in MM9-YEG compared to TSB-D. Accordingly, no strains had biofilm-specific decreases in CFU as calculated as the ratio of biofilm to planktonic growth relative to OG1RF (Figure 5BD). Despite variability in CFU, morphological differences in biofilms were visible. After 4 hr, OG1RF biofilms grew as single cells with small clumps (Figure 5C). OG1RF_10350-Tn biofilms had fewer individual cells and more small chains than OG1RF. Reduced surface coverage was observed in Δ*bph* and OG1RF_10576-Tn biofilms, and OG1RF_10576-Tn biofilms had long chains of cells relative to OG1RF. *prsA*-Tn, *tig*-Tn, and OG1RF_11456-Tn biofilms all had large aggregates of cells. After 24 hr, OG1RF formed dense, thick biofilms with visible cellular aggregates (Figure 5E). Biofilms formed by Δ*bph*, OG1RF_10350-Tn, OG1RF_11456-Tn, OG1RF_11288-Tn had some small aggregates. *prsA*-Tn and *tig*-Tn biofilms had sparse surface coverage with large clusters of cells, and OG1RF_10576-Tn formed biofilms with large aggregates.

**Figure 5.**
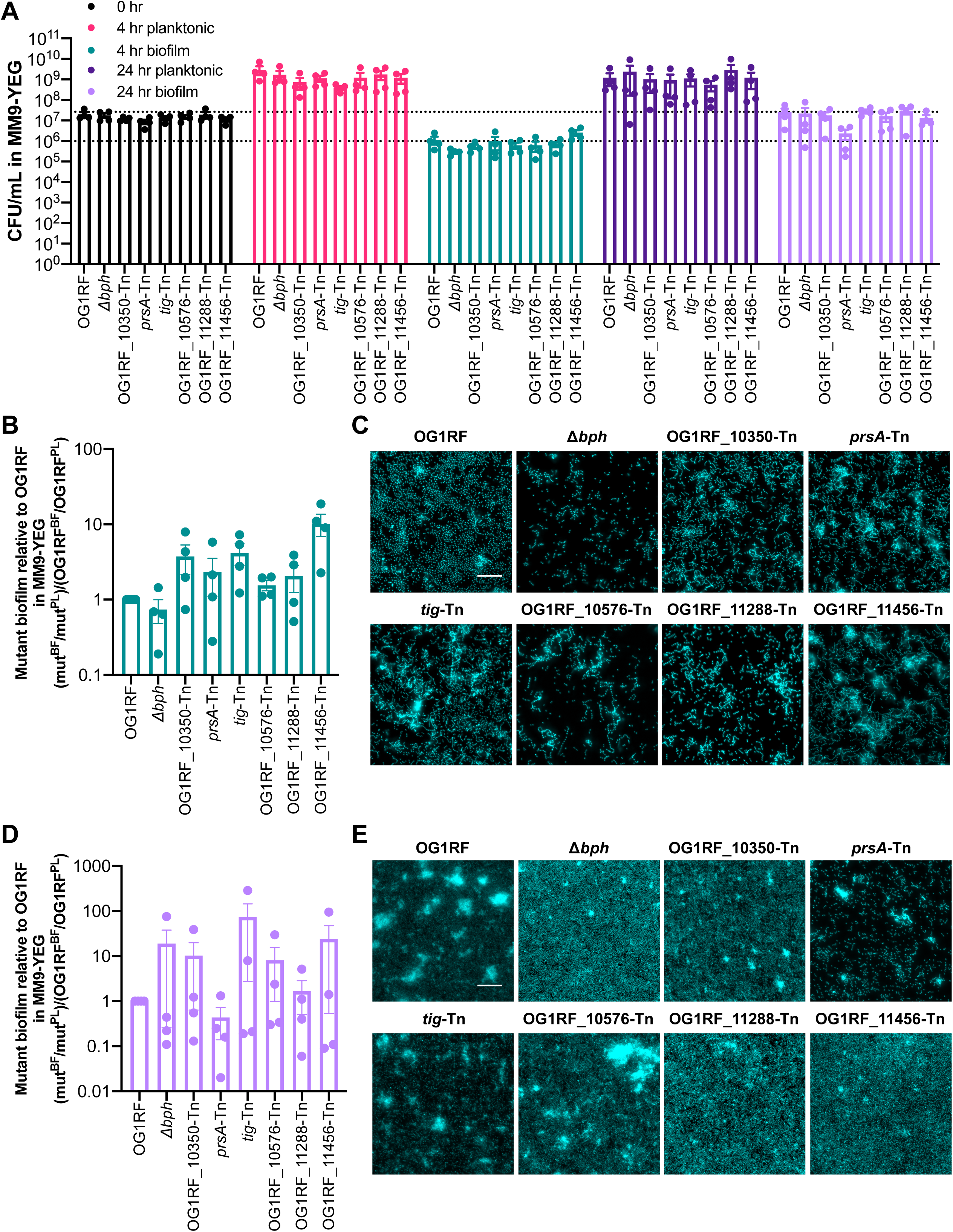
Biofilm formation of selected Tn mutants grown in MultiRep reactors in MM9-YEG. **A)** CFU of strains at 0 hr, 4 hr, and 24 hr. The dotted lines indicated OG1RF biofilm CFU at 24 hr (top line) and 4 hr (bottom line). **B)** Ratio of biofilm to planktonic growth at 4 hr relative to OG1RF. **C)** Representative microscopy images of Hoechst 33342-stained biofilms at 4 hr. **D)** Ratio of biofilm to planktonic growth at 24 hr relative to OG1RF. **E)** Representative microscopy images of Hoechst 33342-stained biofilms at 24 hr. For panel **A**, each data point represents the average of two technical replicates, and a total of four biological replicates were performed. For panels **B** and **D**, data points were derived using the data points shown in panel **A**. Statistical significance was evaluated by one-way ANOVA with Dunnett’s multiple comparisons test. For panels **C** and **E**, samples were grown in parallel to cultures used to generate panel **A**. Scale bars = 20 μm. Two technical replicates were processed for each biological replicates, and representative images are shown.

### Comparative measurements of biofilm growth of OG1RF in different growth assays

Because we observed differences in biofilm morphology depending on growth medium, we used Comstat2 (40) to quantify biomass and thickness of the parental strain using submerged Aclar (6 hr) and MultiRep reactor (4 hr and 24 hr) assays. In general, biofilms grown in MM9-YEG contained more individual cells, whereas biofilms grown in TSB-D had more multicellular chains (Figure 3CF, Figure 4CE, Figure 5CE, Figure S2B). In TSB-D, biomass was not significantly different between submerged Aclar and MultiRep biofilms, nor was biomass of submerged Aclar or 4 hr MultiRep biofilms grown in TSB-D compared to MM9-YEG (Figure S2C). However, the biomass of 24 hr MultiRep biofilms grown in MM9-YEG was 5.3-fold greater than those grown in TSB-D (Figure S2C). In MM9-YEG, 24 hr MultiRep biofilms also had more biomass than 6 hr submerged Aclar biofilms (3.45-fold higher) and 4 hr MultiRep biofilms (13.0-fold higher) (Figure S2C).

We next measured biofilm thickness. Biofilms grown on submerged Aclar for 6 hr or the MultiRep reactor for 4 hr had similar average thicknesses regardless of growth medium (Figure S2D). However, biofilms grown in the MultiRep for 24 hr in MM9-YEG had an average thickness of 23.3 µm, which is 4.06-fold higher than the average thickness of biofilms grown in TSB-D (5.74 µm) and also significantly higher than the other biofilms grown in MM9-YEG (Figure S2D). All biofilms grown in TSB-D had approximately the same maximum thickness (Figure S2E). However, 24 hr MultiRep biofilms grown in MM9-YEG had a maximum thickness of 27.7 µm, which is ∼2-fold more than the other MM9-YEG biofilms and ∼2.5-fold greater than biofilms grown in TSB-D. Taken together, these measurements show that extended cultivation of OG1RF biofilms in MM9-YEG under flow conditions results in thicker biofilms with more biomass than TSB-D, which correlates with the qualitative assessment of biofilm morphology observed using fluorescence microscopy. However, it is currently unknown whether this increase is due solely to the presence of more biofilm cells or to changes in matrix production or composition.

### Tn mutant competition against OG1RF in biofilm co-cultures

The 6 Tn mutants described above were originally identified using TnSeq to evaluate mutant abundance in a community. Therefore, we wanted to measure how the mutants competed in a co-culture with parental OG1RF. In the data reported below, we used both enumeration on selective agar medium (Tn mutants are resistant to chloramphenicol) and fluorescence microscopy to analyze the results of co-cultures. For enumeration, we replaced the Δ*bph* negative control with *bph*-Tn, which has the same biofilm phenotype as the deletion strain (23). To differentially label strains for visualization, we transformed OG1RF with a plasmid expressing tdTomato from a strong constitutive promoter (pP_23_::tdTomato) and each Tn mutant with a plasmid expressing P_23_::GFP. Prior to co-culture, we evaluated whether carriage of the tdTomato or GFP plasmids resulted in growth defects. Two mutants (OG1RF_11456-Tn and OG1RF_11288-Tn) were excluded from co-culture experiments due to poor planktonic growth or unstable fluorescence. With the remaining 4 Tn mutants, we repeated the submerged Aclar experiments described above with cultures in which OG1RF was mixed with single Tn mutants. For all experiments, OG1RF pP_23_::tdTomato was also cultured independently in addition to in co-culture with Tn mutants to ensure that expression of tdTomato did not negatively affect biofilm formation (Figure 6ACEG, Figure 7ACEG, Figure 8ACEG).

For submerged Aclar assays, we inoculated both strains at 10^7^ CFU/mL and quantified OG1RF and Tn mutants after 6 hr. In planktonic cultures grown in TSB-D, only OG1RF-10576-Tn had a significant difference in CFU/mL (∼1 log decrease) relative to OG1RF in the same co-culture (Figure 6B). Biofilm CFU of OG1RF_10576-Tn was also decreased to the same extent relative to OG1RF in co-culture. Interestingly, *prsA*-Tn outgrew OG1RF in these co-culture biofilms by ∼1 log (Figure 6B) and had a 4.23-fold increase in the ratio of biofilm to planktonic CFU relative to OG1RF (Figure S3C), suggesting this mutant outcompeted OG1RF under these conditions. Co-cultures were visualized with fluorescence microscopy (Figure 6CD, Figure S3A). *bph*-Tn/OG1RF biofilms had sparse surface coverage compared to OG1RF alone. The OG1RF_10350-Tn/OG1RF biofilm resembled biofilms formed by the individual strains grown in monoculture. In accordance with CFU quantification, the *prsA*-Tn/OG1RF biofilm had more *prsA*-Tn cells and small clumps than OG1RF. In contrast to *tig*-Tn monoculture biofilms, *tig*-Tn formed large clumps in co-culture with OG1RF. OG1RF_10576-Tn biofilms had low surface coverage when cultured alone, yet this mutant formed large clumps when co-cultured with OG1RF. Interestingly, these large clusters appeared to co-localize with patches of OG1RF cells (Figure S3A).

**Figure 6.**
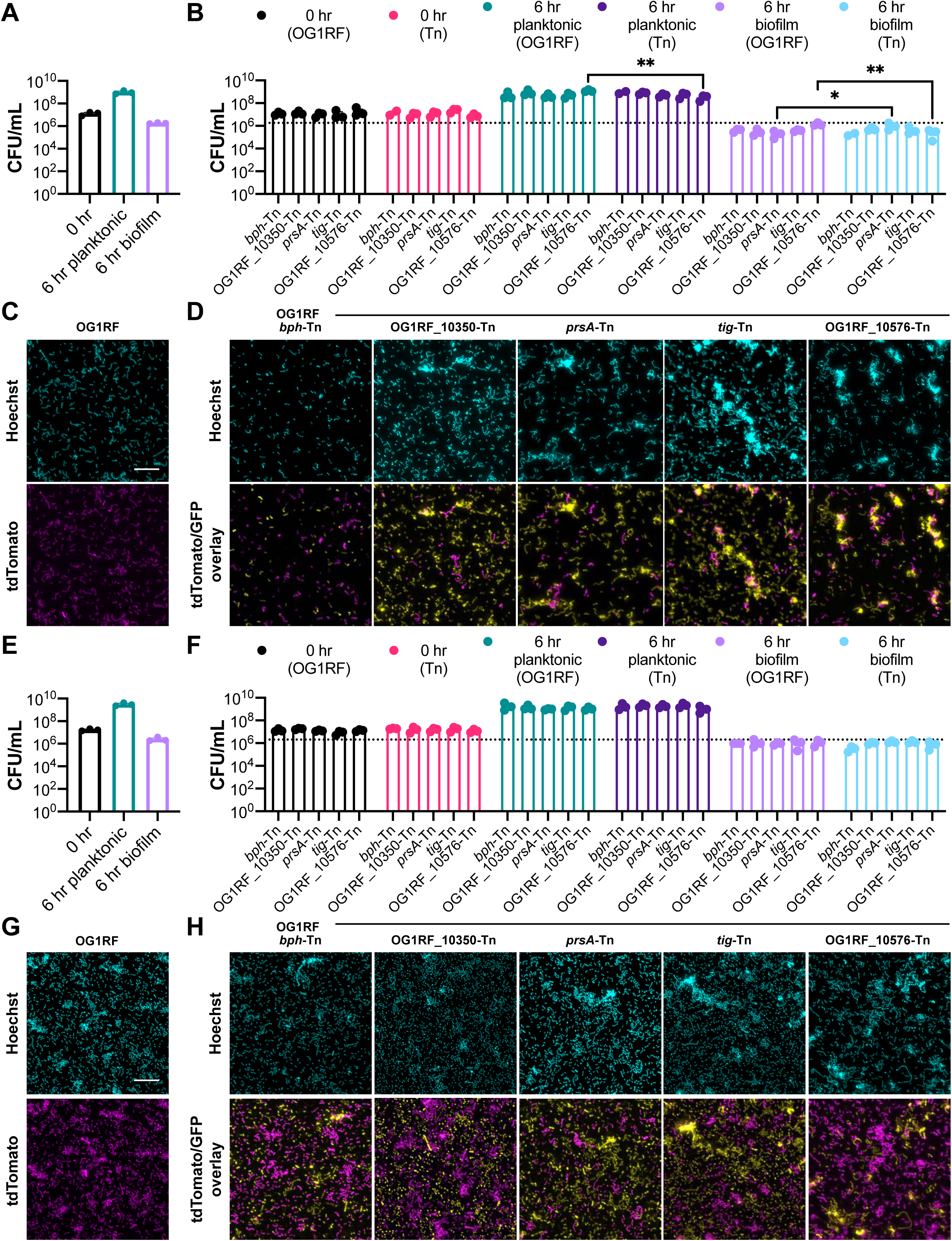
Co-cultures of OG1RF and Tn mutants using the submerged Aclar assay. **A)** CFU of OG1RF grown in TSB-D at 0 hr and 6 hr. **B)** CFU of OG1RF/Tn co-cultures grown in TSB-D at 0 hr and 6 hr. The dotted line indicates biofilm CFU of OG1RF grown in monoculture (value taken from panel **A**). **C)** Representative microscopy images of Hoechst 33342-stained OG1RF pP_23_::tdTomato biofilms grown in TSB-D at 6 hr. **D)** Representative microscopy images of Hoechst 33342-stained OG1RF pP_23_::tdTomato/Tn mutant pP_23_::GFP biofilms grown in TSB-D at 6 hr. **E)** CFU of OG1RF grown in MM9-YEG at 0 hr and 6 hr. **F)** CFU of OG1RF/Tn co-cultures grown in MM9-YEG at 0 hr and 6 hr. The dotted line indicates biofilm CFU of OG1RF grown in monoculture (value taken from panel **E**). **G)** Representative microscopy images of Hoechst 33342-stained OG1RF pP_23_::tdTomato biofilms grown in MM9-YEG at 6 hr. **H)** Representative microscopy images of Hoechst 33342-stained OG1RF pP_23_::tdTomato/Tn mutant pP_23_::GFP biofilms grown in MM9-YEG at 6 hr. For panels **ABEF**, each data point represents the average of two technical replicates, and a total of four biological replicates were performed. Statistical significance was evaluated by two-way ANOVA with Sidak’s multiple comparisons test (*p<0.05, **p<0.01, ***p<0.001, ****p<0.0001). For panels **CDGH**, samples were grown in parallel to cultures used to generate panel **ABEF**. Scale bars = 20 μm. Two technical replicates were processed for each biological replicates, and representative images are shown.

In MM9-YEG, none of the mutants had significantly different planktonic or biofilm CFU relative to OG1RF (Figure 6F). Overall, the MM9-YEG biofilms had more surface coverage than the TSB-D biofilms (compare Figure 6CD and Figure 6GH), and all strains had higher biofilm CFU in MM9-YEG compared to TSB-D. The *bph*-Tn/OG1RF and OG1RF_10350-Tn/OG1RF biofilms resembled those of the mutants and OG1RF grown individually (Figure 6GH, Figure S3B). However, *prsA*-Tn, *tig*-Tn, and OG1RF_10576-Tn biofilms contained fewer aggregates when co-cultured with OG1RF than when grown individually. Additionally, biofilms from *tig*-Tn co-cultured with OG1RF in MM9-YEG contained more individual *tig*-Tn cells (as opposed to multicellular chains) than when co-cultured in TSB-D. OG1RF_10576-Tn formed clumps and chains with visibly less surface coverage than OG1RF (Figure S3B) when co-cultured with OG1RF in MM9-YEG, although there was no statistical difference between OG1RF_10576-Tn and OG1RF biofilm CFU.

### Biofilm formation of OG1RF and Tn mutant co-cultures in miniature flow reactors

Biofilm formation of co-cultures was evaluated using the MultiRep biofilm flow chambers described above, and each strain was inoculated at 10^7^ CFU/mL. After 4 hr in TSB-D, there were no statistically significant differences in planktonic CFU between OG1RF and any mutants, but OG1RF_10350-Tn/OG1RF biofilms contained 3.6-fold more OG1RF_10350-Tn CFU than OG1RF CFU (Figure 7B). Visualization of biofilms revealed that OG1RF_10350 and *tig*-Tn formed biofilms with aggregates containing both mutant and OG1RF cells (Figure 7D, Figure S4A). *prsA*-Tn formed large aggregates in co-culture with OG1RF, but these aggregates contained relatively few OG1RF cells (Figure 7D, Figure S4A). *bph*-Tn/OG1RF and OG1RF_10576-Tn/OG1RF biofilms had less surface coverage than OG1RF grown alone (Figure 7CD). After 24 hr growth in TSB-D, there were no significant differences in co-culture planktonic or biofilm CFU (Figure 7F). OG1RF pP_23_::tdTomato and co-culture biofilms grew as monolayers of mostly individual cells, with fewer multicellular aggregates and less chaining than observed after 4 hr (Figure 7GH). Fewer *bph*-tn and OG1RF_10576-Tn were present relative to OG1RF (Figure 7H, Figure S4B), although only *bph*-Tn had significantly reduced biofilm CFU relative to OG1RF (Figure 7F).

**Figure 7.**
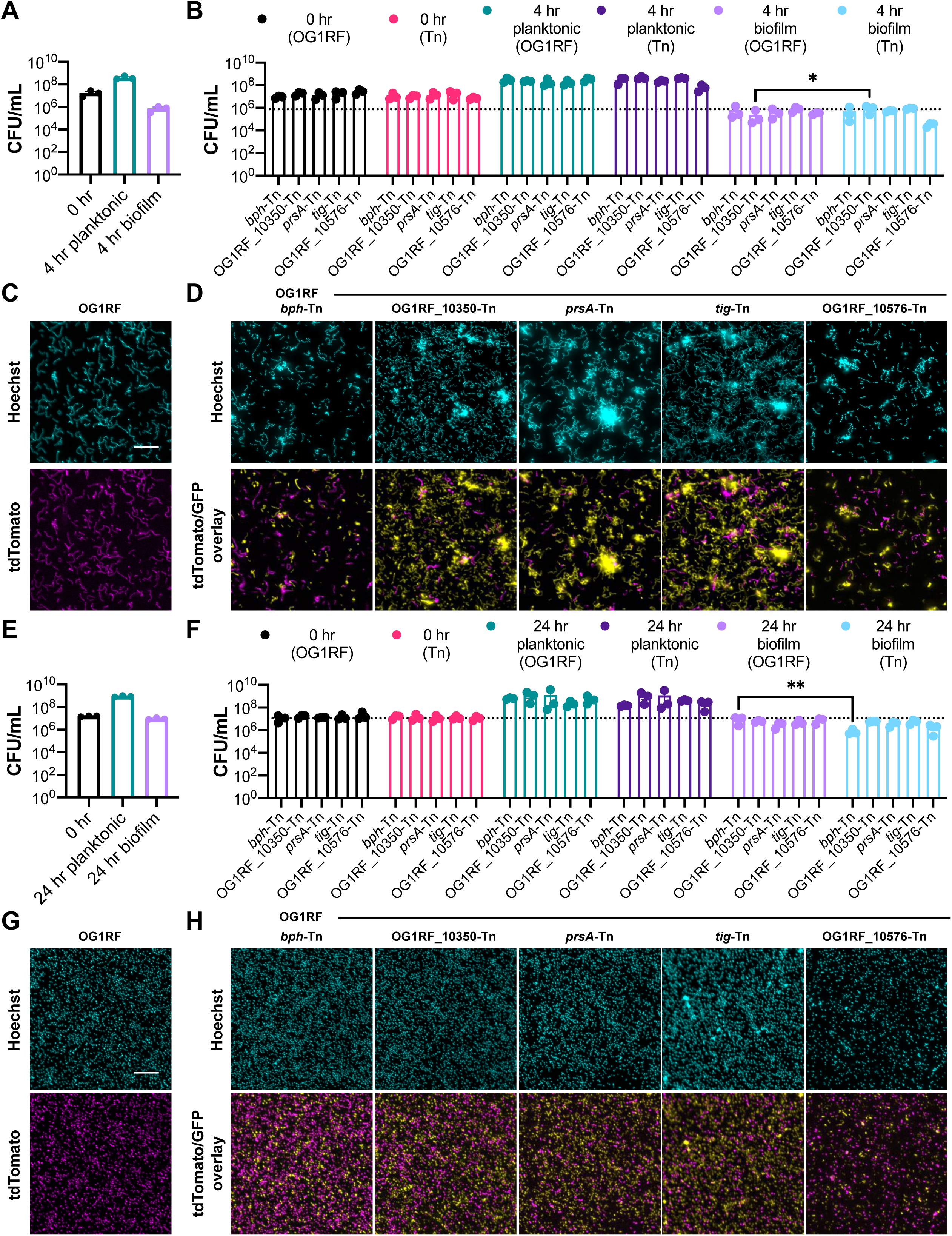
Co-cultures of OG1RF and Tn mutants in TSB-D in the MultiRep reactors. **A)** CFU of OG1RF at 0 hr and 4 hr. **B)** CFU of OG1RF/Tn co-cultures at 0 hr and 4 hr. The dotted line indicates biofilm CFU of OG1RF grown in monoculture (value taken from panel **A**). **C)** Representative microscopy images of Hoechst 33342-stained OG1RF pP_23_::tdTomato biofilms at 4 hr. **D)** Representative microscopy images of Hoechst 33342-stained OG1RF pP_23_::tdTomato/Tn mutant pP_23_::GFP biofilms at 4 hr. **E)** CFU of OG1RF at 0 hr and 24 hr. **F)** CFU of OG1RF/Tn co-cultures at 0 hr and 24 hr. The dotted line indicates biofilm CFU of OG1RF grown in monoculture (value taken from panel **E**). **G)** Representative microscopy images of Hoechst 33342-stained OG1RF pP_23_::tdTomato biofilms at 24 hr. **H)** Representative microscopy images of Hoechst 33342-stained OG1RF pP_23_::tdTomato/Tn mutant pP_23_::GFP biofilms at 24 hr. For panels **ABEF**, each data point represents the average of two technical replicates, and a total of three biological replicates were performed. Statistical significance was evaluated by two-way ANOVA with Sidak’s multiple comparisons test (*p<0.05, **p<0.01, ***p<0.001, ****p<0.0001). For panels **CDGH**, samples were grown in parallel to cultures used to generate panel **ABEF**. Scale bars = 20 μm. Two technical replicates were processed for each biological replicates, and representative images are shown.

After 4 hr in MM9-YEG, there were no significant differences in planktonic or biofilm CFU between OG1RF and Tn mutants in co-culture (Figure 8B). Very few mutant cells were visible in the *bph*-Tn/OG1RF and OG1RF_10576-Tn/OG1RF biofilms (Figure 8D, Figure S5A). OG1RF_10350/OG1RF biofilms had larger aggregates of cells than those grown in TSB-D for 4 hr. *prsA*-Tn/OG1RF and *tig*-Tn/OG1RF biofilms resembled those grown in TSB-D for 4 hr and contained large aggregates of cells. After 24 hr growth in MM9-YEG, there were no significant differences between OG1RF or Tn mutant CFUs in planktonic or biofilm cultures (Figure 8F). Co-culture biofilms contained thick multicellular aggregates of both OG1RF and Tn mutants, with the exception of *prsA*-Tn co-culture biofilms, which had fewer large aggregates (Figure 8H, Figure S5B). None of the Tn mutants had significant differences in the ratio of biofilm to planktonic cells relative to OG1RF at either 4 hr or 24 hr (Figure S5CD).

**Figure 8.**
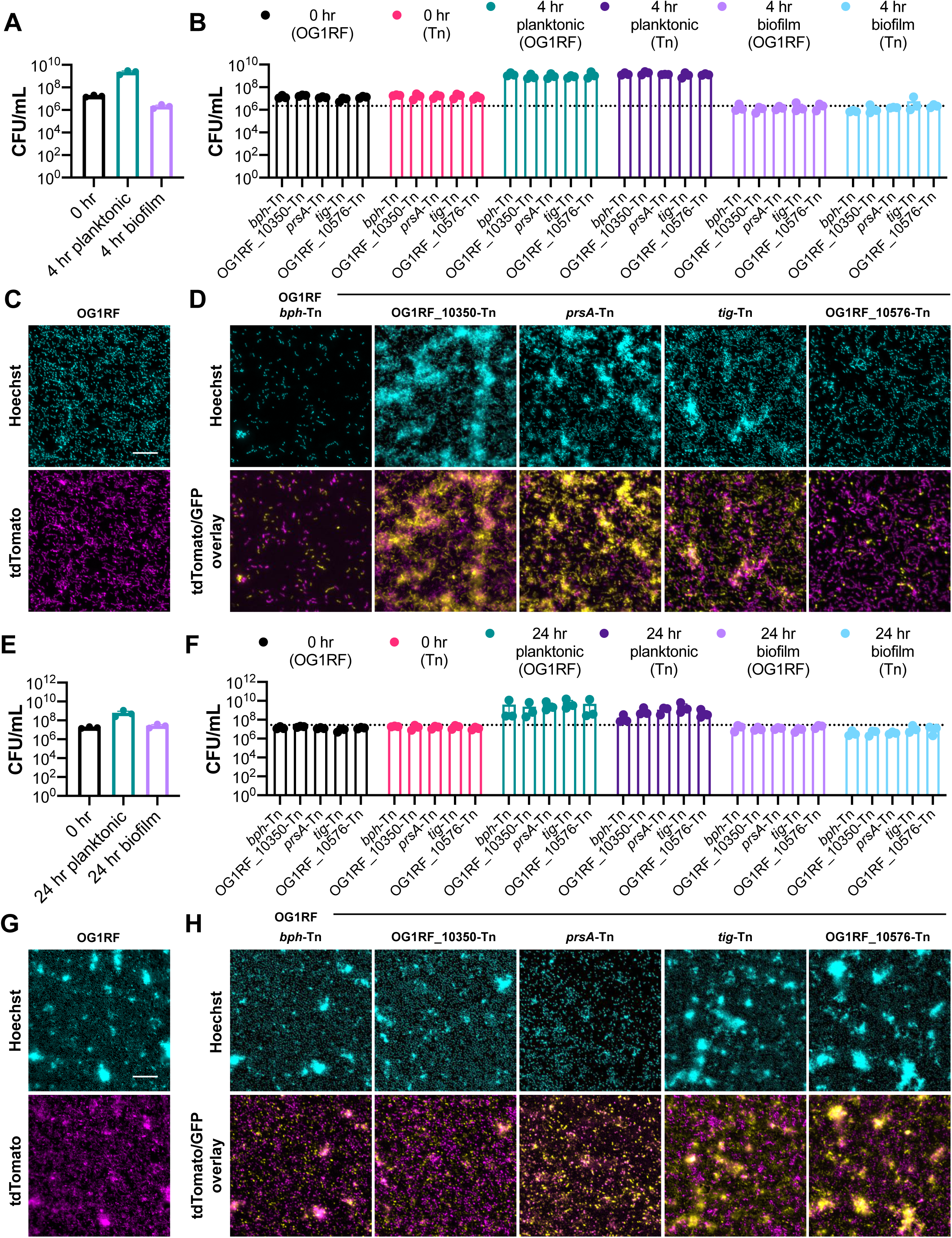
Co-cultures of OG1RF and Tn mutants in MM9-YEG in the MultiRep reactors. **A)** CFU of OG1RF at 0 hr and 4 hr. **B)** CFU of OG1RF/Tn co-cultures at 0 hr and 4 hr. The dotted line indicates biofilm CFU of OG1RF grown in monoculture (value taken from panel **A**). **C)** Representative microscopy images of Hoechst 33342-stained OG1RF pP_23_::tdTomato biofilms at 4 hr. **D)** Representative microscopy images of Hoechst 33342-stained OG1RF pP_23_::tdTomato/Tn mutant pP_23_::GFP biofilms at 4 hr. **E)** CFU of OG1RF at 0 hr and 24 hr. **F)** CFU of OG1RF/Tn co-cultures at 0 hr and 24 hr. The dotted line indicates biofilm CFU of OG1RF grown in monoculture (value taken from panel **E**). **G)** Representative microscopy images of Hoechst 33342-stained OG1RF pP_23_::tdTomato biofilms at 24 hr. **H)** Representative microscopy images of Hoechst 33342-stained OG1RF pP_23_::tdTomato/Tn mutant pP_23_::GFP biofilms at 24 hr. For panels **ABEF**, each data point represents the average of two technical replicates, and a total of three biological replicates were performed. Statistical significance was evaluated by two-way ANOVA with Sidak’s multiple comparisons test (*p<0.05, **p<0.01, ***p<0.001, ****p<0.0001). For panels **CDGH**, samples were grown in parallel to cultures used to generate panel **ABEF**. Scale bars = 20 μm. Two technical replicates were processed for each biological replicates, and representative images are shown.

### Putative biochemical activities of newly identified biofilm determinants from structural modeling and a functional assay

Between 10-40% of bacterial gene products are poorly characterized or annotated as hypothetical (41), although they are frequently identified as loci of interest in experiments in OG1RF and other organisms (22, 29, 42, 43). Of the 45 new genes identified as biofilm determinants from TnSeq (Table 3), 6 were annotated as hypothetical, as gene products that are incongruous with known *E. faecalis* biology (chemotaxis or sporulation), or had conflicting annotations across multiple databases (NCBI and KEGG). Others had vague annotations, and their function had not been studied in *Enterococcus*. We used Phyre2 (44) to predict structures for 14 proteins for which we tested the corresponding Tn mutants in 96-well plate biofilm assays (Table S3), including 3 chosen for analysis with microscopy and co-cultures (OG1RF_10350-Tn, OG1RF_11288-Tn, and OG1RF_11456). OG1RF_10350 and OG1RF_11288 are annotated in different databases as LytR-Cps2a-Psr (LCP)-family proteins or transcriptional regulators. Early studies on LCP-family proteins suggested they could be transcription factors, but the well-characterized examples are phosphotransferases that catalyze attachment of glycopolymers to the cell wall of Gram-positive bacteria (45). OG1RF_10350 and OG1RF_11288 have only 25.08% sequence homology but are predicted to have similar core crystal structures with distal helices encompassing putative transmembrane domains (Figure S6A). Predicted structural homologs of these proteins included putative transcription factors and uncharacterized proteins but also well-characterized cell wall modifying enzymes such as Csp2A from *Streptococcus pneumoniae* D39 (PDB 4DE8 (46)), LcpA from *Staphylococcus aureus* N315 (PDB 6UEX (47), and TagU from *Bacillus subtilis* 168 (PDB 6UF6 (47)) (Table S3, Figure S6B). This suggests that OG1RF_10350 and OG1RF_11288 may modify the *E. faecalis* cell wall, which could affect the ability of these mutants to form biofilms under the conditions we tested.

OG1RF_11456 is annotated as a methyl-accepting chemotaxis receptor, although *E. faecalis* is non-motile. Biofilms formed by OG1RF_11456-Tn contained large multicellular aggregates (Figure 3CF, Figure 4C, Figure 5C). Phyre2 analysis of OG1RF_11456 yielded high confidence matches to the methylation and signaling domains of Tsr, the membrane-bound serine chemotaxis receptor from *E. coli* (PDB 1QU7 (48)), and Tm14, a chemoreceptor from *Thermatoga maritima* (PDB 3G67 (49)) (Table S3). The putative structure of OG1RF_11456 is an extended linear conformation, similar to Tsr and Tm14 (Figure S6C). OG1RF_11456 has a predicted transmembrane domain that best aligns with the Tsr/Tm14 signaling domains, which are cytoplasmic (48, 49). Although the Tsr methylation sites are not conserved in OG1RF_11456, this protein contains multiple glutamine and glutamic acid residues that could be involved in signal transduction. However, additional experiments are needed to confirm whether OG1RF_11456 functions as a signaling protein in *E. faecalis* and how this relates to the extreme clumping phenotypes observed in OG1RF_11456-Tn biofilms.

Numerous *in vitro* biofilm determinants of OG1RF have also been characterized as virulence factors in models of biofilm-associated infections (5). One such protein is GelE (gelatinase), a secreted metalloprotease regulated by the Fsr quorum sensing system; *gelE^-^* mutants show defects in biofilm formation in vitro and are attenuated in animal models (50, 51). Therefore, we tested whether the 43 Tn mutants chosen for 96-well plate biofilm assays could secrete active GelE. Mutants were spotted on agar plates containing 3% gelatin, and colonies were evaluated for production of an opaque zone indicative of gelatinase activity (51). All mutants except for *prsA*-Tn (OG1RF_10423-Tn) had gelatinase-positive phenotypes similar to OG1RF (Figure S6). PrsA is a predicted extracellular membrane-bound peptidyl-prolyl cis-trans isomerase that is associated with tolerance to salt stress (52), *E. faecalis* virulence in *Galleria mellonella* (52), and is upregulated in a rabbit subdermal abscess model (42), although no specific protein substrates for chaperone or foldase activity have been identified. We suspect that PrsA enhances correct folding of GelE as it transits the membrane during secretion. The cumulative results from this study suggest important roles for several poorly characterized gene products as important modulators of biofilm formation and architecture.

## Discussion

In this study, we cultured a library of *E. faecalis* OG1RF Tn mutants in CDC biofilm reactors and identified new determinants of biofilm formation using TnSeq. We identified core biofilm determinants in OG1RF by comparing our results to previous studies done using microtiter plate biofilm assays (23, 27). While the endpoint measurement of both experiments is biofilm formation, microtiter plate assays test the ability of a strain to form a biofilm when grown as a monoculture, whereas TnSeq measures fitness of a community of mutants. As such, it is expected that some mutants behave differently in these assays, and there is value in using TnSeq to study biofilm formation even in species or strains that have been extensively used in microtiter plate experiments. Using the same Tn library to identify biofilm determinants in multiple conditions can allow for categorization of core biofilm determinants and condition-specific accessory determinants. Core biofilm determinants could be promising targets for the development of new anti-biofilm or antimicrobial therapeutics.

We used 2 growth media (TSB-D and MM9-YEG) to generate a more comprehensive view of how growth conditions affect *E. faecalis* biofilms. These results demonstrate that growth medium can significantly influence genetic determinants of biofilm formation, given the number of mutants identified in TSB-D compared to MM9-YEG as well as the small overlap of mutants identified in both media. Additionally, an increase in multicellular chains was observed in TSB-D biofilms compared to those grown in MM9-YEG (Figure S2B, and compare Figure 4C with Figure 5C), whereas OG1RF biofilms grown in MM9-YEG for 24 hr were thicker than those grown in TSB-D. Glucose availability is a significant difference between TSB-D (no added glucose) and MM9-YEG (0.4% added glucose), although other nutritional differences could affect biofilm formation. This provides rationale for testing multiple growth conditions during genetic screens and suggests that nutritional availability in different host niches, such as the GI tract compared to wounds or abscesses, could affect determinants of biofilm growth.

Examining temporal biofilm formation also revealed important morphological variations. In general, biofilms cultured for 24 hr in the MultiRep reactors had a marked decrease in cell chain length compared to biofilms cultured for 4 hr. However, multiple factors such as time or fluid flow could influence these architectural changes. Based on our results, extrapolating the influence of biofilm determinants between growth conditions should be done with caution; previously we found that only a minority of genes identified as biofilm determinants using in vitro screens affected virulence in experimental infections involving biofilm growth (34). Additional work is needed to understand how nutrient availability and the temporal nature of biofilm development affects biofilm determinants, biofilm morphology, and matrix composition at different sites of infection or colonization, including niches not associated with a mammalian host.

Validating mutants identified in a primary screen is a major challenge with TnSeq and other high-throughput genetic experiments. Here, we tested biofilm-deficient mutants identified from CBR TnSeq in three subsequent biofilm assays (microtiter plates, submerged Aclar, and MultiRep reactors) that represent a tradeoff between throughput and similarity to the primary screen. Microtiter plate assays allow simultaneous testing of dozens to hundreds of mutants using small sample volumes, but they are “closed” systems incubated under static conditions without supplementation of fresh growth medium. Despite the dissimilarity of microtiter plates and CBRs, ∼30% of the Tn mutants we tested had defects in biofilm formation in 96-well plates, suggesting that these may be a reasonable platform for secondary screens of large sets of mutants in order to identify those with reproducible phenotypes for subsequent studies. However, this must be balanced against the probability of excluding mutants with CBR-specific (or flow-specific) biofilm-deficient phenotypes. Although submerged Aclar assays and MultiRep reactors can more closely mimic the conditions of CBRs, these are more suitable for smaller sets of mutants given the time and resources required to process, quantify, and visualize samples. Fresh growth medium can be provided to cultures grown in the MultiRep reactors, enabling the study of biofilms under flow conditions with lower reagent requirements than CBRs and increasing feasibility of studies in the presence of antibiotics or other compounds.

From the underrepresented Tn mutants identified in biofilm TnSeq, we chose 6 mutants for quantification and visualization of biofilms. Importantly, quantification of biofilm cells did not correlate with biofilm morphology. Relying on quantitative measurements of biofilm formation to identify differences between strains may obscure important variances in morphology or developmental processes such as biofilm remodeling or cellular exodus (16). Quantification of biofilm and planktonic cells also suggested that the Tn mutants used in co-cultures could compete with OG1RF under most conditions. Interestingly, we found that *prsA*-Tn grew better when co-cultured with OG1RF in TSB-D than when grown alone. However, these Tn mutants were originally identified as underrepresented in TnSeq, so perhaps the complexity of the Tn library restricts growth of certain mutants in biofilms. Of the 4 genes encoding proteins with peptidyl-prolyl isomerase (PPIase) domains in OG1RF (*prsA*/EF0685, *tig*/EF0715, OG1RF_11253/EF1534, and OG1RF_12199/EF2898 (52)), only *prsA*-Tn and *tig*-Tn mutants were underrepresented in our biofilm study. Disruptions in both genes led to altered biofilm morphology relative to OG1RF, with mutant biofilms containing large aggregates of cells. Additionally, *prsA*-Tn had a gelatinase-negative phenotype when grown on gelatin plates, but *tig*-Tn was gelatinase-positive. Determining the substrates of the OG1RF PPIases is crucial for understanding how aberrant protein folding and secretion affect biofilm architecture and growth.

Multiple genes in the *epa* operon were also underrepresented in biofilm TnSeq. With the exception of *epaQ*, these are all part of the variable region downstream of genes encoding the core rhamnopolysaccharide backbone (53). Modification of the Epa backbone or side chains affects biofilm architecture, antibiotic-associated biofilm formation, and resistance to phage and antibiotics (16, 25, 26, 32, 33, 53, 54). However, our previous studies on EpaOX and EpaQ did not identify them as important for biofilm formation in the absence of antibiotics or cell wall stressors (26, 33). These studies quantified biofilm formation in microtiter plates, so perhaps these *epa* genes are important for biofilm integrity in the presence of shear stress generated in CBRs. Recently, Guerardel *et al*. proposed that addition of teichoic acid to the rhamnan backbone and anchoring of Epa to the cell wall may be mediated by LCP-family proteins (53). OG1RF encodes 5 LCP-family proteins, 2 of which we identified as important for biofilm formation (OG1RF_10350 and OG1RF_11288). The predicted crystal structures of these proteins have high homology to LCP-family wall teichoic acid transferases in other Gram-positive bacteria (45, 46). Interestingly, OG1RF_10350-Tn and OG1RF_11288-Tn biofilms had increased chaining and clumping relative to OG1RF when grown in MM9-YEG, and *epaOX* and *epaQ* mutant strains also form biofilms with altered morphology (16, 33). Additional work is needed to identify the targets and substrates of LCP-family proteins in OG1RF and how cell wall integrity and composition is affected in their absence.

Overall, our study identified sets of new and core biofilm determinants for *E. faecalis* OG1RF, and that disruption of multiple biofilm determinants leads to drastic changes in biofilm morphology during monoculture and co-culture. We also identified specific morphological signatures of OG1RF biofilms grown in different media, with biofilms grown in TSB-D containing mostly multicellular chains and biofilms grown in MM9-YEG containing mostly single cells. Many newly identified biofilm determinants are poorly characterized proteins or intergenic regions, suggesting that our understanding of enterococcal biofilm formation in diverse conditions is still incomplete. Additionally, we identified potential roles in production of gelatinase and Epa or cell wall homeostasis for multiple new biofilm determinants. Taken together, our work shows how *E. faecalis* biofilm architecture can be modified by growth medium, experimental conditions, and genetic determinants, demonstrating that comparing biofilms across multiple conditions can provide new insights into the process of biofilm formation as well as basic bacterial biology.

## Materials and Methods

### Bacterial strains and growth conditions

Bacterial strains were maintained as freezer stocks at −80 °C in 20-25% glycerol. Strains were routinely grown in brain-heart infusion (BHI) broth for cloning and generating freezer stocks. All strains used in this study are listed in Table S4. Overnight cultures were grown in the same medium used for experiments. Antibiotics were used at the following concentrations: chloramphenicol (Cm) 10 µg/mL, erythromycin (Erm) 10 µg/mL (*E. faecalis*) or 80 µg/mL (*E. coli*), fusidic acid (FA) 25 µg/mL, tetracycline (Tet) 5 (liquid) or 10 (plates) µg/mL. When required, agar was added to growth medium at a final concentration of 1% (w/v). MM9-YEG (modified M9 growth medium supplemented with yeast extract and glucose) was prepared as previously described (28). BHI and tryptic soy broth without added dextrose (TSB-D) were purchased from BD and prepared according to manufacturer’s instructions. Fusidic acid was purchased from Chem-Impex, and all other antibiotics were purchased from Sigma.

### Cloning and Tn mutant verification

Nucleotide sequences of primers are listed in Table S3. All restriction enzymes were purchased from New England Biolabs. For construction of the cCF10-inducible *tig* complementation vector, *tig* was amplified from purified OG1RF genomic DNA using Pfu Ultra II polymerase (Agilent), digested with BamHI-HF/NheI-HF, and ligated to pCIEtm (23) treated with the same restriction enzymes. The plasmid construct was verified by Sanger sequencing (Eurofins). For generation of constitutive fluorescent protein constructs, P_23_ was excised from pDL278p23 (55) by digesting with EcoRI-HF/BamHI-HF and ligated to pTCV-LacSpec digested with the same restriction enzymes. A fragment encoding promoterless GFP (56) flanked by BamHI and BlpI sites was inserted to create pP_23_::GFP, and the BamHI-SphI fragment from pJ201::187931 was inserted to create pP_23_::tdTomato. The Tn insertions in strains used for submerged Aclar and MultiRep reactor experiments were verified by colony PCR using the oligos listed in Table S4. The Tn insertion adds ∼2.1 kb to the size of the wild-type allele.

### CDC biofilm reactors

Reactors were assembled as previously described (14, 16) and incubated at 37 °C overnight to ensure a lack of contamination. Polycarbonate (BioSurfaces Technologies Corp.) and Aclar (Electron Microscopy Sciences) coupons were used as biofilm substrates. Immediately prior to inoculation, single-use Tn library aliquots were removed from storage at −80 °C and thawed on ice. Growth medium (either MM9-YEG or TSB-D) was inoculated with 6 x 10^8^ – 2 x 10^9^ CFU. Batch cultures were grown without flow for 4-6 hours after which the peristaltic pump (Cole Parmer) was turned on at a flow rate of 8 mL/minute for 18-20 hours (total experiment time = 24 hours). Two biological replicate reactors were run for each Tn library/growth medium combination.

### DNA isolation, library preparation, and transposon sequencing

Substrates were removed from the CDC biofilm reactor chamber and processed to remove adherent biofilm cells. Polycarbonate coupons were aseptically removed and placed in 6-well plates (4 coupons/well) containing 5 mL distilled water and incubated for 5 min at room temperature to remove non-adherent cells. To obtain attached biofilm cells, 12 coupons were placed in 50 mL conical tubes containing 30 mL KPBS (potassium phosphate-buffered saline, pH 7.0) with 2 mM EDTA and vortexed in a Benchmixer multi-tube vortexer (Benchmark Scientific) at 2,000 rpm for 5 minutes. Biofilms grown on Aclar membranes were rinsed in 50 mL conical tubes with 30 mL KPBS followed by inversion to remove non-adherent cells. Rinsed Aclar were submerged in 4 mL KPBS with 2 mM EDTA, and biofilms were removed by scraping with a sterile razor blade. Biofilms from multiple substrates from each reactor were pooled in a conical tube, pelleted at 6371× g for 15 min, and frozen at −80 °C until further use. Pellets were resuspended in 180 uL enzymatic lysis buffer (20 mM Tris-HCl pH 8.0, 2 mM EDTA, 1.2% Triton X-100) with 30 mg/mL lysozyme and 500 U/mL mutanolysin. After 30 min incubation at 37 °C, 25 uL Proteinase K and 200 uL Buffer AL (DNeasy Blood and Tissue Kit, Qiagen) were added. Tubes were incubated at 55 °C for 30 min, after which DNA was extracted using a DNeasy Blood and Tissue Kit following manufacturer’s instructions. Samples were submitted to the University of Minnesota Genomics Center for library preparation and sequencing. Sequencing libraries were prepared using the Illumina TruSeq Nano library preparation kit as previously described (22). Libraries were sequenced as 125-bp paired-end reads on an Illumina HiSeq 2500 in high output mode (440M reads total).

Sequencing reads were processed using a published workflow (22). Briefly, reads were trimmed and aligned to the OG1RF genome (NC_017316.1), and Tn insertions at TA sites were quantified. Statistical significance of the relative abundance of Tn reads at each TA site was evaluated using a chi-squared test and an additional Monte Carlo-based method. Scripts for all processing steps are publicly available (https://github.com/dunnylabumn/Ef_OG1RF_TnSeq). Output files were filtered for nucleotide positions of Tn mutants known to be present in the library based on previous sequencing (22). Log_2_ fold changes were calculated from relative Tn abundances. Statistical significance was defined as p < 0.05 and a Monte Carlo simulation value of 1.119552, the lowest value obtained in these calculations.

Tn mutants used for additional experiments were obtained from frozen library stock plates and grown on BHI/FA agar plates. Single colonies were picked and patched onto BHI/Erm to ensure loss of the plasmids used in Tn mutagenesis and BHI/Cm to confirm functionality of the Cm resistance gene located in the Tn. Single colonies were picked from BHI/Cm plates and grown in BHI/Cm/FA to generate freezer stocks. Tn insertions were verified by colony PCR using primers flanking the gene of interest (Table S3). The Tn adds ∼2.1 kb to the size of the parental allele (27, 57).

### Scanning electron microscopy (SEM)

Biofilms were removed from the CBRs and rinsed with KPBS three times, then processed for SEM using the cationic dye stabilization methods described previously (14–16, 33). Briefly, biofilms were subjected to primary fixation in sodium cacodylate buffer containing methanol-free EM-grade formaldehyde (2%), glutaraldehyde (2%), sucrose (4%), and alcian blue 8GX (0.15%) overnight. Coupons were then rinsed 3x with sodium cacodylate buffer and subjected to secondary fixation in sodium cacodylate buffer containing 1% osmium tetroxide and 1.5% potassium ferrocyanide for 1 hr. Fixed samples were rinsed 3x with sodium cacodylate buffer and chemically dried using a graded ethanol series, processed in a CO_2_-based critical point dryer (Tousimis, Rockville, MD), and sputter coated with ∼2 nm iridium (EM ACE600; Leica, Buffalo Grove, IL). Imaging was done using a Hitachi SU8230 field emission instrument at 0.8 kV using the low-angle backscatter and secondary electron detectors.

### Biofilm assays

96-well plate biofilm assays were carried out as described previously (23, 26, 29). Overnight cultures for complementation assays were grown with 5 µg/mL tetracycline and 25 ng/mL cCF10, and experiments were performed in the indicated growth medium supplemented with 25 ng/mL cCF10. Briefly, overnight cultures were diluted 1:100 in the appropriate growth medium, and 100 µL was added to a 96-well plate (Corning 3935). For the secondary screens using 43 Tn mutants, two technical replicates were performed for each strain. For complementation assays, three technical replicates were performed for each strain. For all experiments, values shown are the results of three independent biological replicates. Plates were incubated in a humidified plastic container at 37 °C for the indicated amount of time. Cell growth was measured in a Biotek Synergy HT plate reader as the absorbance at 600 nm (A_600_). Plates were gently washed three times with ultrapure water using a Biotek plate washer, dried in a biosafety cabinet or on a lab bench overnight, and stained with 100 µL 0.1% safranin (Sigma). Stained plates were washed three times and dried. A_450_ was measured to quantify safranin-stained biofilm biomass. Biofilm production was evaluated as the ratio of stained biofilm biomass to overall growth (A_450_/A_600_), and values were normalized to biofilm production of OG1RF.

For submerged Aclar biofilm assays, overnight cultures were adjusted to 10^7^ CFU/mL in the appropriate growth medium, and 1 mL was added to 1 well of a 24-well plate (Costar 3524) with a 5 mm Aclar disc. Plates were incubated at 37 °C in a plastic container on a tabletop shaker (Thermo Scientific MaxQ 2000) at 100 rpm. After 6 hr, planktonic cells were transferred to microfuge tubes. Aclar discs were washed by gently shaking in KPBS and transferred to microfuge tubes with 1 mL KPBS (1 Aclar/tube). Tubes with planktonic cultures and Aclar discs were vortexed at 2500 rpm for 5 min in a Benchmixer multi-tube vortexer (Benchmark Scientific), then diluted (10-fold serial dilutions) in KPBS and plated on BHI/FA medium to enumerate colonies. For co-culture experiments, diluted cultures were plated on BHI/FA (total CFU counts) and BHI/Cm plates (Tn mutant CFU counts). CFU/mL values for OG1RF in co-culture were obtained by subtracting the CFU/mL counts from BHI/Cm plates from the CFU/mL counts from BHI/FA plates. At least three biological replicates (each with two technical replicates) were performed for all strains.

MultiRep biofilm reactors (Stratix Labs, Maple Grove, MN) were loaded with 5 mm Aclar discs (6 Aclar per channel). Influx (MasterFlex HV-96117-13) and efflux (MasterFlex EW-06424-16) tubing was attached to each channel and capped with foil prior to autoclaving. The 10% growth medium was autoclaved in a separate bottle with sterile connecting tubing and attached to the influx reactor tubing immediately prior to inoculation. Overnight cultures were diluted to 1 x 10^7^ CFU/mL, and 4 mL was added to each channel (1 channel per strain). The reactor was sealed by placing 2 silicon sheets in the lid and clamping the lid on the reactor using Irwin Quick-Grip ratcheting bar clamps. The influx tubing was connected to peristaltic pumps (MasterFlex 77202-60), and the efflux tubing was placed horizontally over waste containers. Reactors were kept at 37 °C with static incubation for 4 hr, after which the pumps were turned on at a flow rate of 0.1 mL/min for 20 hr. For disassembly and sample processing, the reactor lids were removed, and 2 mL planktonic culture was transferred to microfuge tubes. Aclar were removed and rinsed in KPBS, then placed in microfuge tubes with 1 mL KPBS. Tubes with planktonic cultures and Aclar were vortexed, diluted, and enumerated as described above.

### Fluorescence microscopy

For all experiments, Aclar coupons (2 per strain) were rinsed 3 times in KPBS and stained for 15 min in Hanks’ Balanced Salt Solution with CaCl_2_ and MgCl_2_ (Gibco) and 5 µg/mL Hoechst 33342 (Molecular Probes) with gentle agitation. After staining, Aclar were washed 3 times in fresh KPBS and transferred to a 48-well plate (Costar 3548) with 1 mL 10% buffered formalin (Fisher Scientific) with gentle agitation and shielded from light for 12-16 hours. After fixing, Aclar were washed in KPBS and mounted on a Superfrost Plus microscope slide (Fisher Scientific) in a 0.24 mm double-sided adhesive Secureseal spacer (Grace BioLabs) with a 7 mm hole punched to accommodate the Aclar. Aclar were covered with 7 uL Prolong Glass Antifade Mountant (Invitrogen) and a Gold Seal cover slip (#1.5, Fisher Scientific). Slides were cured at room temperature shielded from light for 4-8 hours and stored at 4 °C until imaging.

### Microscopy and image processing

All images were acquired on a Zeiss Axio Imager M1 widefield microscope with a Plan-APO 20× (0.8 numerical aperture (NA)) using an X-Cite 120 metal halide light source (EXFO, Inc.) illuminating 365 nm, 470 nm, and 550 nm excitation filters for Hoechst 33342, GFP, or tdTomato, respectively. Images were captured using the Zeiss AxioCam 503 mono microscope camera and Zen imaging software (v 2.1, Zeiss). For each Aclar coupon, two independent images were obtained, yielding four images per sample from which a final representative image was chosen. Representative images were processed using the Fiji ImageJ package (version 1.48v; NIH) and subjected to background subtraction with a rolling ball radius of 50 pixels using the internal ImageJ function as well as uniformly applied brightness and contrast adjustments of the entire image prior to cropping (58). For biofilm co-culture images, the Hoechst, GFP, and tdTomato images were false colored cyan, yellow, and magenta (respectively) using Fiji. For co-culture images, tdTomato (OG1RF) and GFP (Tn mutant) maximum intensity projections were processed independently and merged. Images were cropped to 500×500 pixels using GIMP (v 2.0) and exported as PNG files. The GFP (mutant) and tdTomato (OG1RF) MIPs were processed independently and merged.

Biofilm thickness and distribution were analyzed using Comstat2. Cells were imaged using an Axio Observer.Z1 confocal microscope equipped with an LSM 800-based Airyscan system in normal confocal mode (Zeiss). Confocal images were acquired with a 20× 0.8 NA objective and 405-nm lasers for excitation of Hoechst 33342 stain. For image analysis, two representative *z* stacks were taken per Aclar coupon with a 1 μm interval. Each experiment used three independent biological replicates with at least 2 Aclar coupons in each. Maximum thickness of the biofilms was determined from the Hoechst channel using the Comstat2.1 plugin for ImageJ (40, 59). All image processing adheres to the standards outlined by Rossner and Yamada (60).

### Gelatinase assays

Overnight cultures were grown in the respective growth medium and were spotted onto TSB-D agar plates supplemented with 3% gelatin (w/v). Plates were incubated overnight at 37 °C then moved to 4 °C for 1-3 hours prior to imaging. Plate photos were obtained using a ProteinSimple (Cell Biosciences) FluorChem FC3 imager. Strains were considered gelatinase positive if they developed a halo around colony growth and gelatinase negative if no halo was present.

### Bioinformatic analysis

Functional annotations of proteins were obtained from KEGG and NCBI. Protein sequences were obtained from NCBI and used as input for Phyre2 in intensive mode (44). Transmembrane predictions were done using TMHMM (61). Additional protein structure files were downloaded from PDB, and structures were rendered in Pymol 2.1 (62).

### Statistical analysis

All statistical analysis was carried out using GraphPad Prism (version 9.0.1). Statistical tests and significance are described in the figure legends. Corrections for multiple comparisons were performed using the test recommended by GraphPad.

## Acknowledgments

We thank the University of Minnesota Genomics Center for assistance with TnSeq and the Minnesota Supercomputing Institute (MSI) at the University of Minnesota for providing computational resources. Parts of this work were carried out in the Characterization Facility, University of Minnesota, which receives partial support from the NSF through the MRSEC (Award Number DMR-2011401) and the NNCI (Award Number ECCS-2025124) programs. This work was supported by 1RO1AI122742 to G. M. D. from the NIH. J. L. E. W. was supported by American Heart Association Grant #19POST34450124 / Julia Willett / 2018. M. L. K. was supported by grants TL1R002493 and UL1TR002494 from the NIH’s National Center for Advancing Translational Sciences. A. M. T. B. received support via NIH training grant AI055433 for portions of this work. We thank Dr. Elizabeth Cameron for providing the pCIEtm::*tig* plasmid and Dawn Manias for assistance with constructing pP_23_::GFP and pP_23_::tdTomato.

---

**Figure S1.**
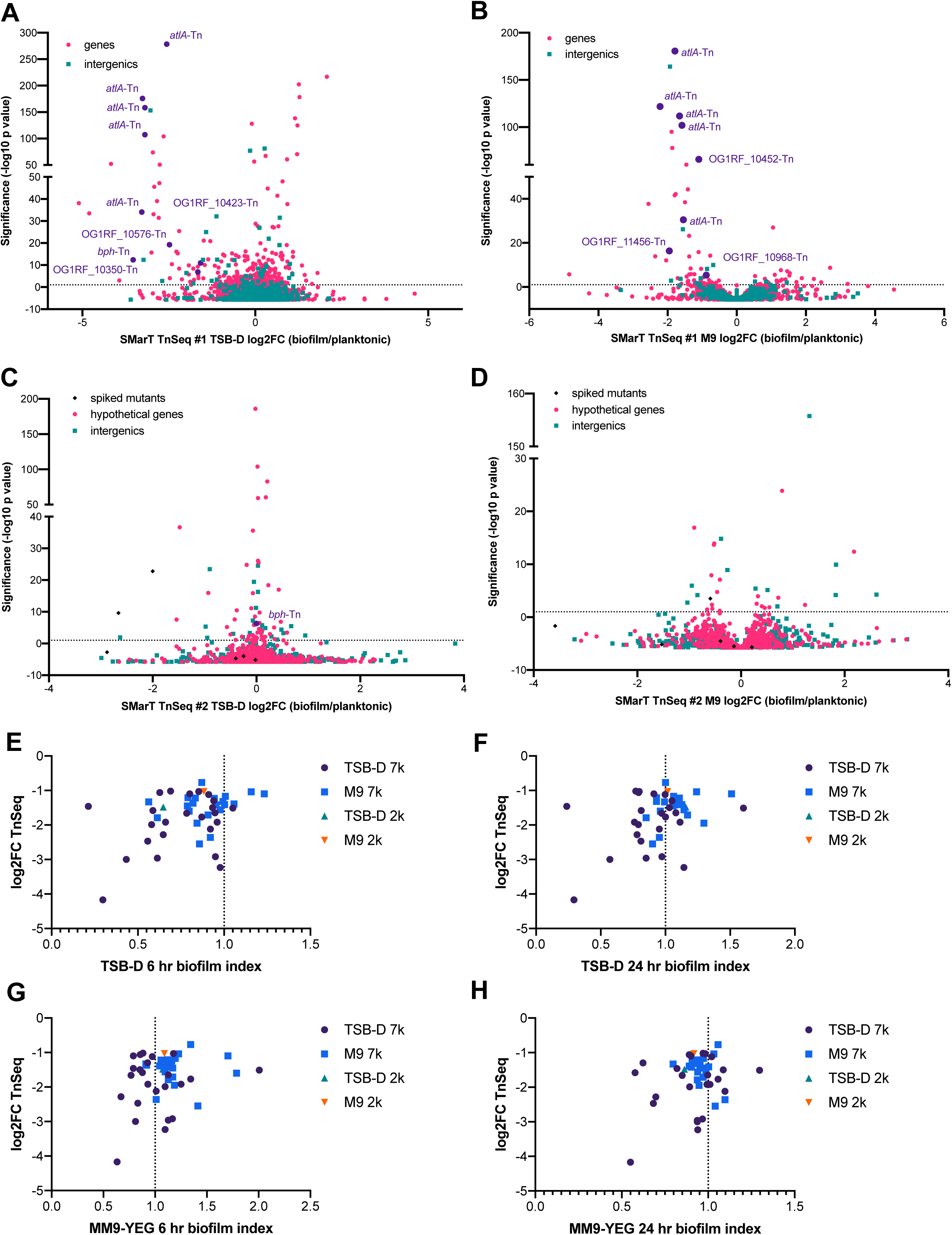
Relative abundance of Tn mutants in CBR TnSeq and comparison with biofilm formation in microtiter plates. Panels **A-D** show data from CBR TnSeq, and panels **E-H** compare the fitness of mutants selected from the TnSeq to their phenotypes in monocultures using microtiter plate biofilm assays. Volcano plots of SmarT TnSeq library #1 (6,829 mutants) in **A)** TSB-D and **B)** MM9-YEG and SmarT TnSeq library #2 (1,948 mutants) in **C)** TSB-D and **D)** MM9-YEG. Tn mutants previously identified as biofilm determinants or chosen for microtiter plate assays are highlighted in purple. Log_2_FC values from biofilm TnSeq were compared to biofilm index values obtained from microtiter plate biofilm assays for **E)** 6 hr biofilms in TSB-D, **F)** 24 hr biofilms in TSB-D, **G)** 6 hr biofilms in MM9-YEG, and **H)** 24 hr biofilms in MM9-YEG.

**Figure S2.**
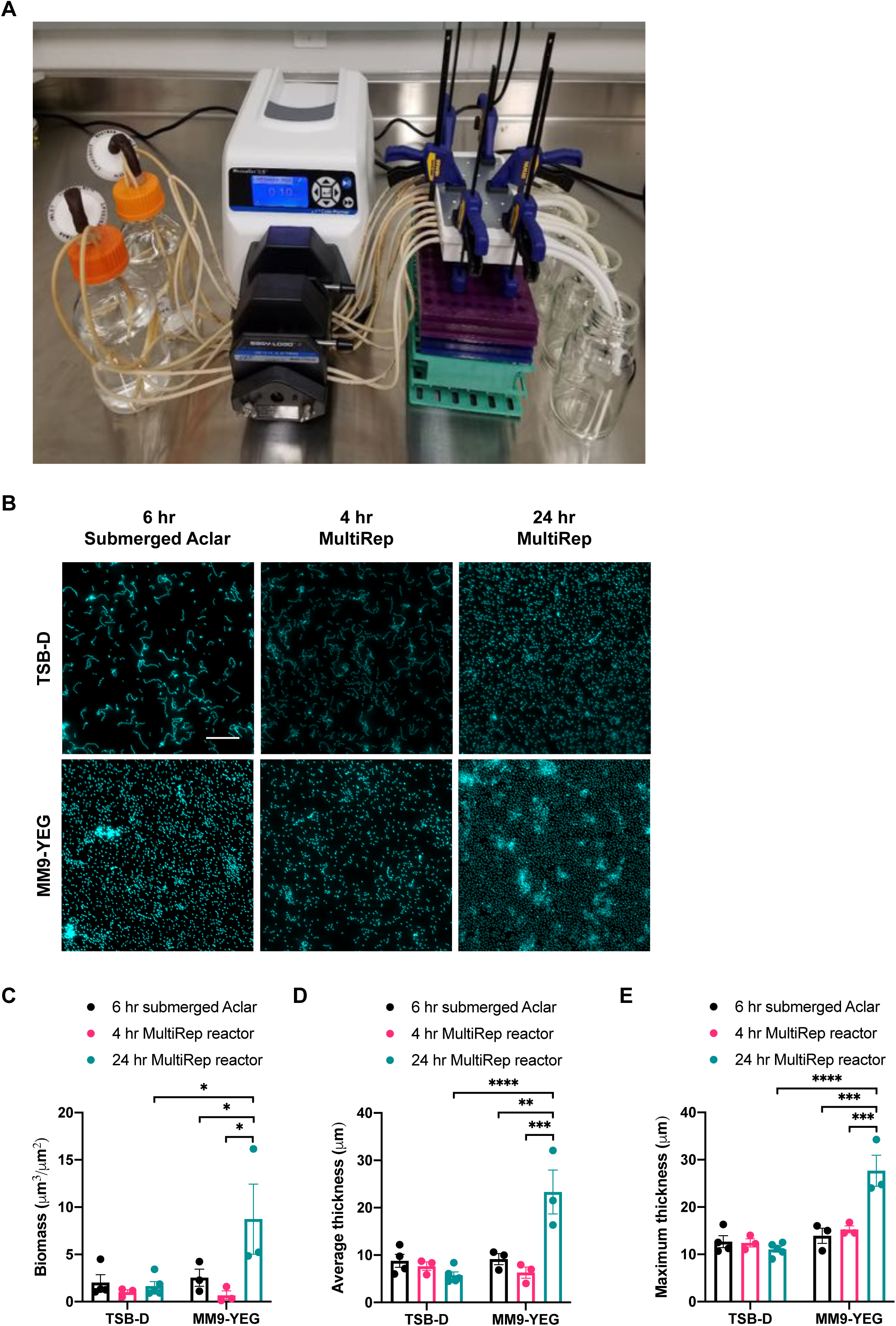
MultiRep biofilm reactors and analysis of OG1RF biofilms grown under multiple experimental conditions. **A)** Photograph showing an assembled MultiRep biofilm reactor. Bottles with sterile growth medium are shown on the left, and outflow tubes with waste containers are shown on the right. **B)** Additional fluorescence microscopy images of OG1RF biofilms obtained during biological replicates of experiments shown in Figure 3, Figure 4, and Figure 5. Scale bars = 20 μm. Images of OG1RF biofilms were used for Comstat2 analysis of **C)** overall biomass, **D)** average biofilm thickness, and **E)** maximum biofilm thickness. Statistical significance was evaluated by two-way ANOVA with Tukey’s multiple comparisons test (*p<0.05, **p<0.01, ***p<0.001, ****p<0.0001).

**Figure S3.**
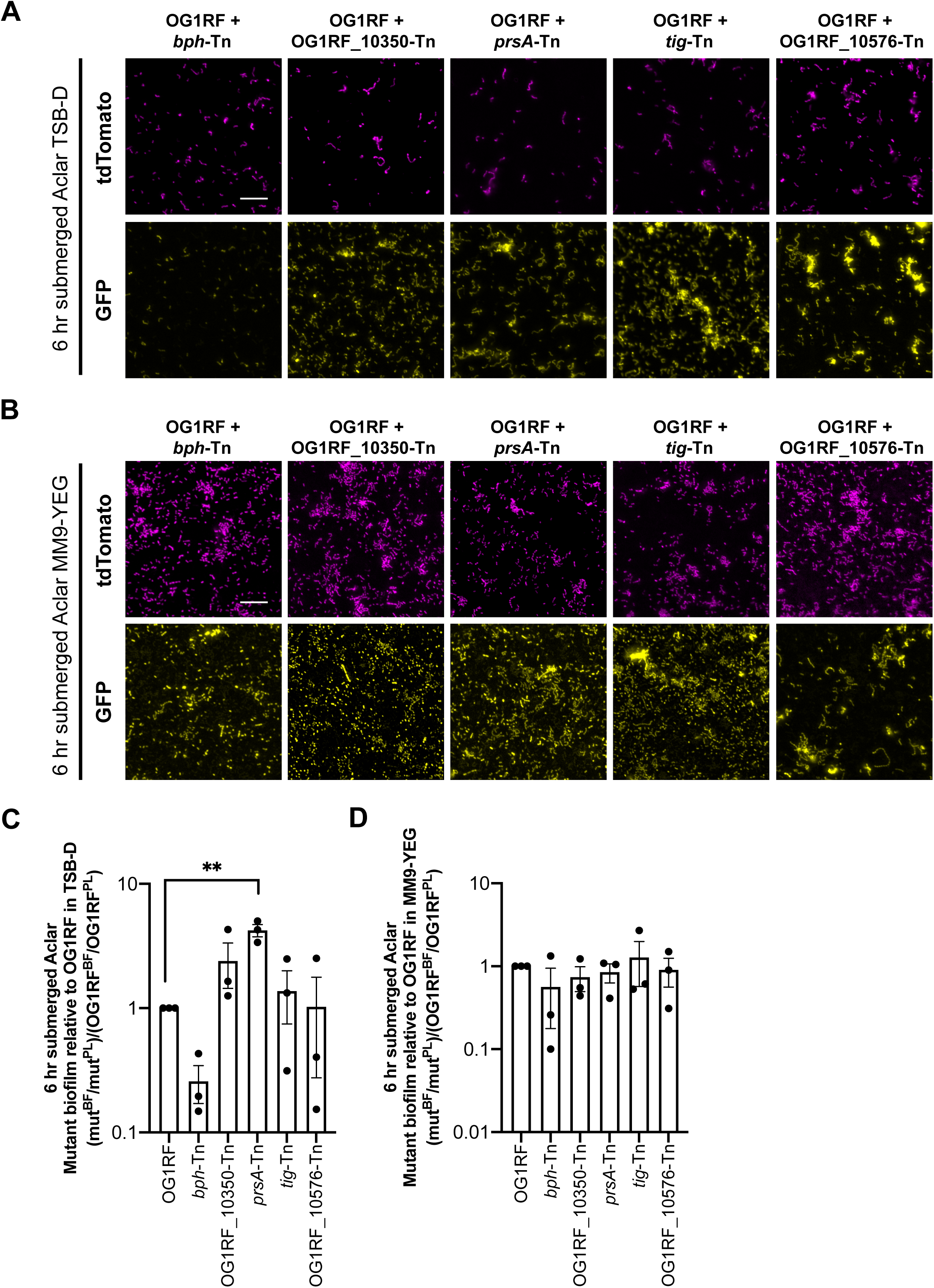
Individual channels and relative biofilm growth of Tn mutants in submerged Aclar co-cultures. The individual tdTomato and GFP panels for **A)** TSB-D co-cultures and **B)** MM9-YEG co-cultures that are shown as overlays in Figure 6DH are presented here for clarity. Scale bars = 20 μm. The ratio of biofilm to planktonic growth relative to OG1RF were calculated for **C)** TSB-D co-cultures and **D)** MM9-YEG co-cultures. Data points in **C** and **D** were calculated from the CFU values presented in Figure 6. Statistical significance was evaluated by two-way ANOVA with Dunnett’s multiple comparisons test (*p<0.05, **p<0.01, ***p<0.001, ****p<0.0001).

**Figure S4.**
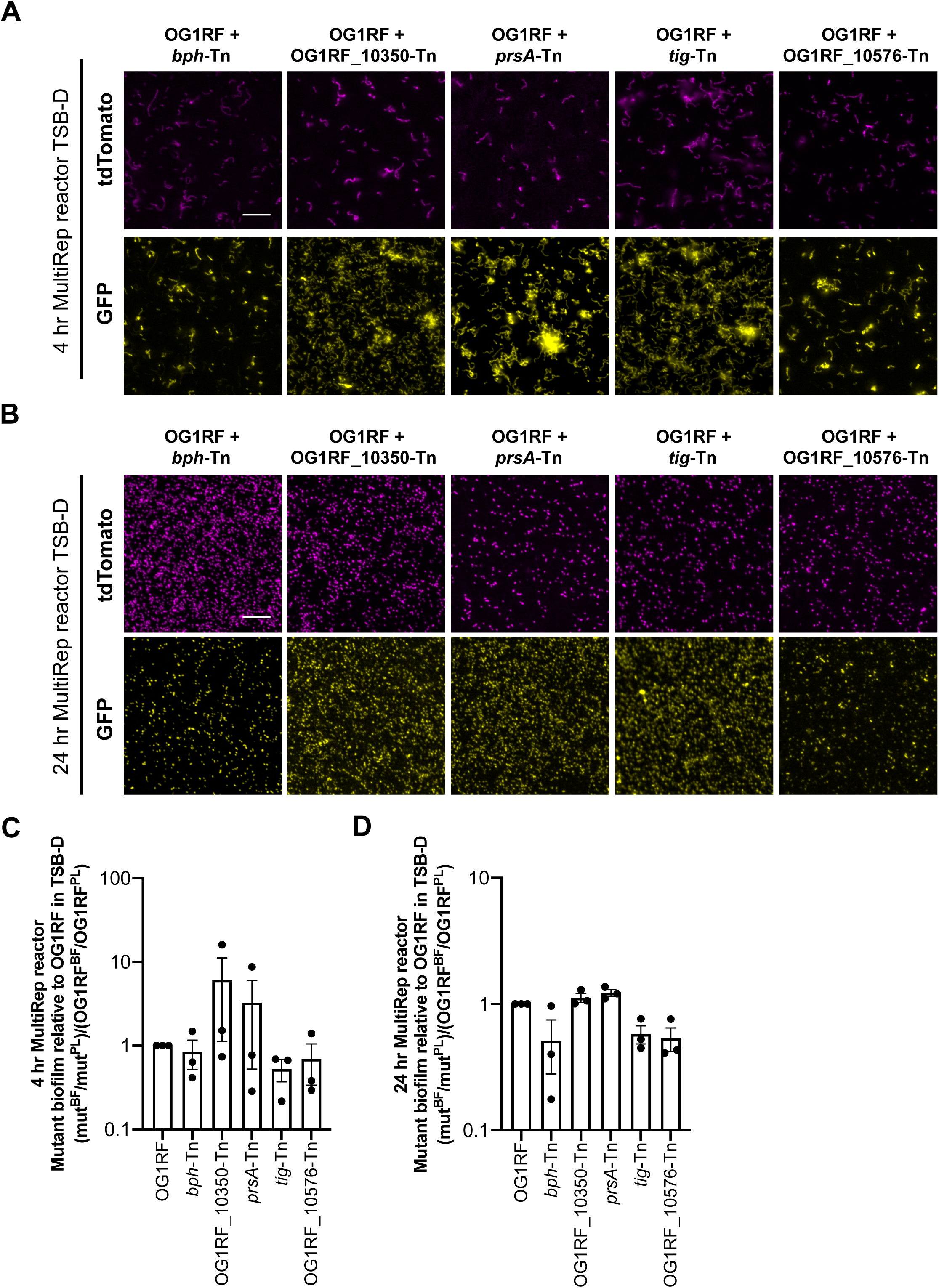
Individual channels and relative biofilm growth of Tn mutants in TSB-D in the MultiRep reactors. The individual tdTomato and GFP panels for **A)** 4 hr and **B)** 24 hr co-cultures that are shown as overlays in Figure 7DH are presented here for clarity. Scale bars = 20 μm. The ratio of biofilm to planktonic growth relative to OG1RF were calculated for **C)** 4 hr and **D)** 24 hr co-cultures. Data points in **C** and **D** were calculated from the CFU values presented in Figure 7. Statistical significance was evaluated by two-way ANOVA with Dunnett’s multiple comparisons test (*p<0.05, **p<0.01, ***p<0.001, ****p<0.0001).

**Figure S5.**
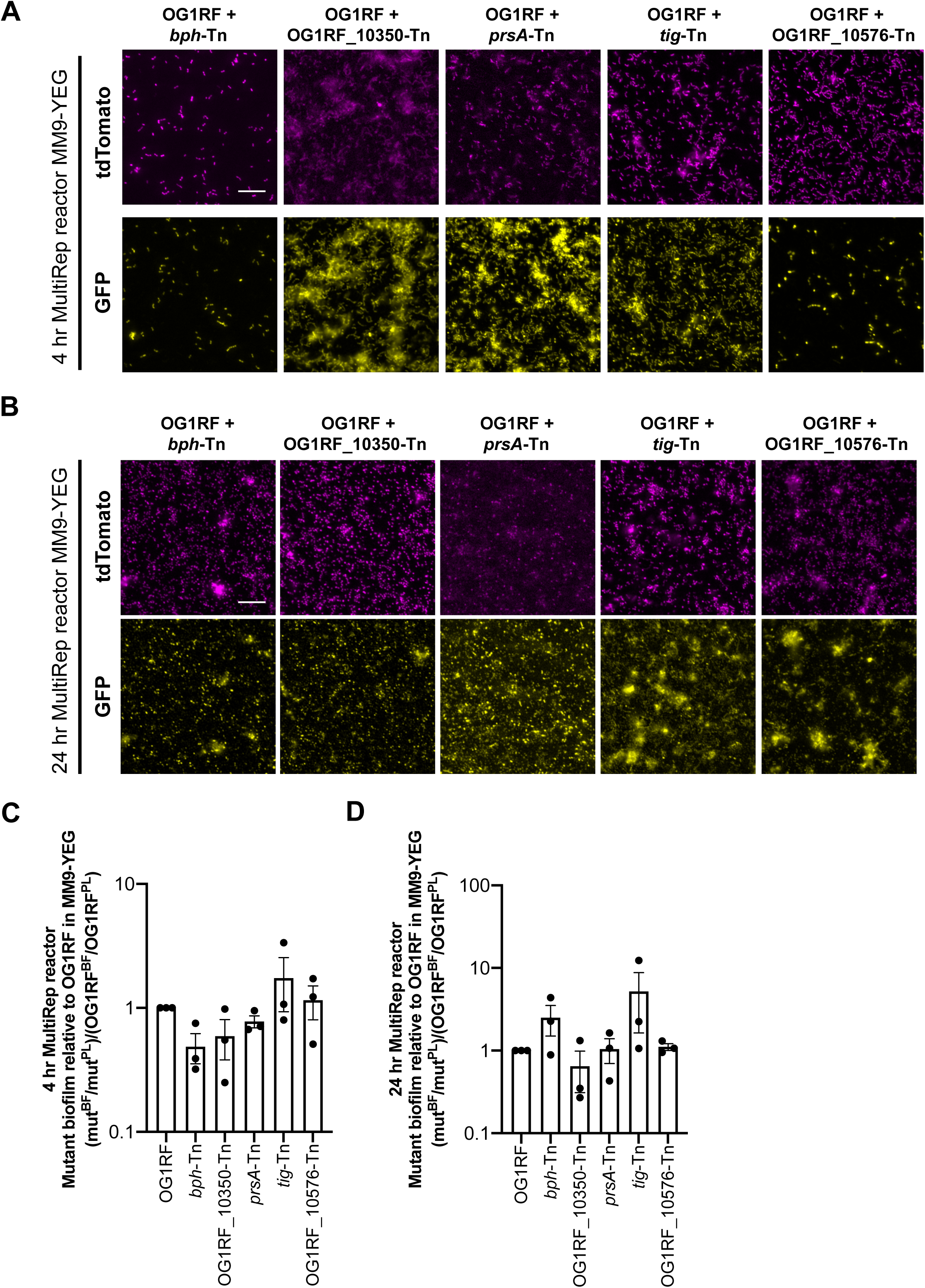
Individual channels and relative biofilm growth of Tn mutants in MM9-YEG in the MultiRep reactors. The individual tdTomato and GFP panels for **A)** 4 hr and **B)** 24 hr co-cultures that are shown as overlays in Figure 8DH are presented here for clarity. Scale bars = 20 μm. The ratio of biofilm to planktonic growth relative to OG1RF were calculated for **C)** 4 hr and **D)** 24 hr co-cultures. Data points in **C** and **D** were calculated from the CFU values presented in Figure 8. Statistical significance was evaluated by two-way ANOVA with Dunnett’s multiple comparisons test (*p<0.05, **p<0.01, ***p<0.001, ****p<0.0001).

**Figure S6.**
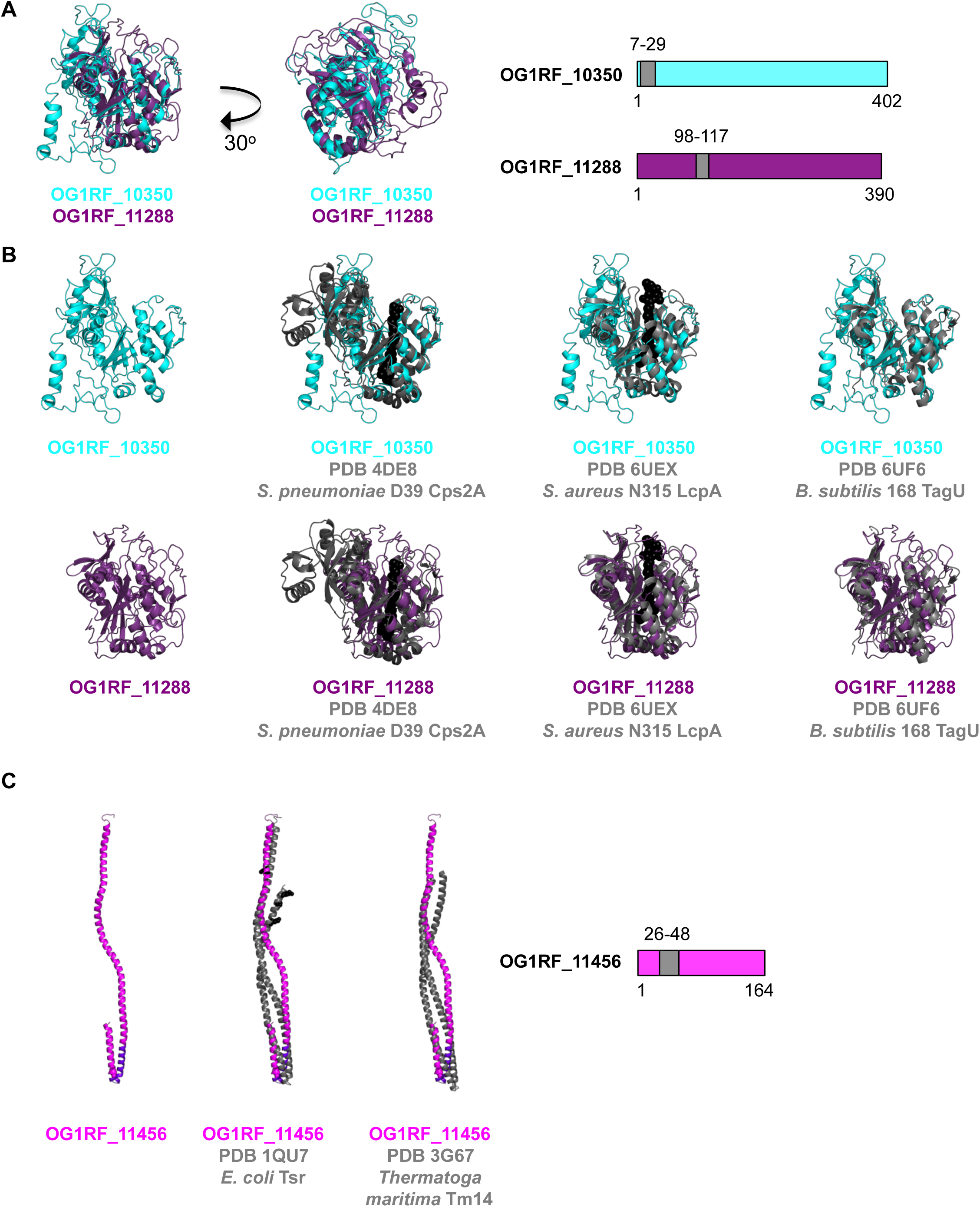
Predicted crystal structures for OG1RF_10350, OG1RF_11288, and OG1RF_11456. **A)** Phyre2 was used to predict the structures of OG1RF_10350 and OG1RF_11288. Both proteins have predicted transmembrane domains (shown as gray boxes in cartoons on the right). **B)** OG1RF_10350 and OG1RF_11288 have predicted structural homology to multiple LCP-family wall teichoic acid transferases from Gram-positive bacteria. PDB identifiers for Cps2A, LcpA, and TagU are shown. Lipid substrates for Cps2A and LcpA are represented as black spheres. **C)** The putative crystal structure of OG1RF_11456 has predicted structural homology to membrane-bound chemosensors Tsr and Tm14. OG1RF_11456 has one predicted transmembrane domain (shown as a gray box in the cartoon on the right and as black residues in the OG1RF_11456 predicted structure). Tsr residues that undergo methylation are shown as black spheres.

**Figure S7.**
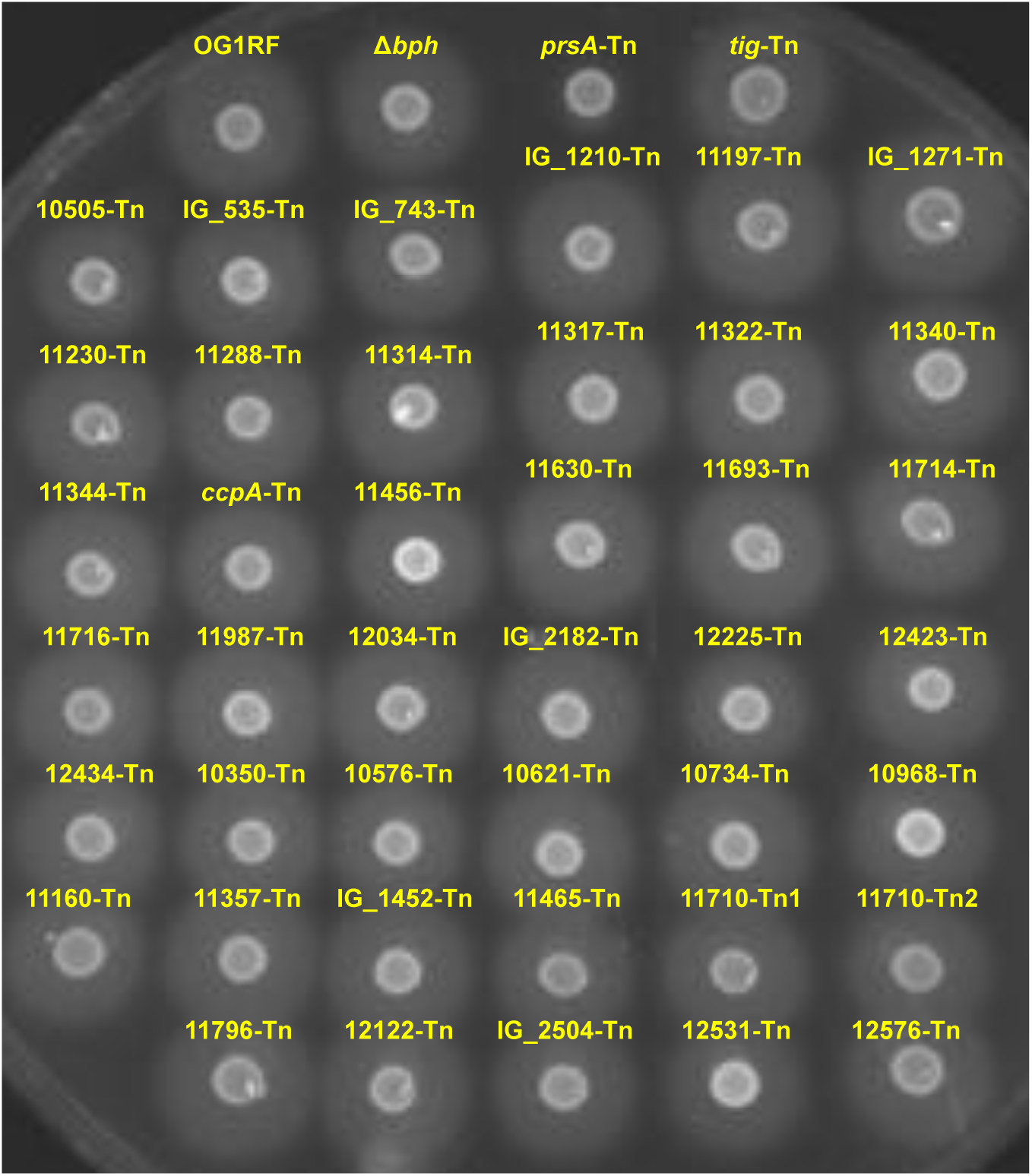
Gelatinase activity of Tn mutants chosen for microtiter plate biofilm assays. Overnight cultures grown in TSB-D were spotted onto a TSB-D agar plate supplemented with 3% gelatin. After overnight growth, plates were refrigerated until the zone surrounding colonies was visible. Three biological replicates were performed, and a representative image is shown.

**Table S3.**
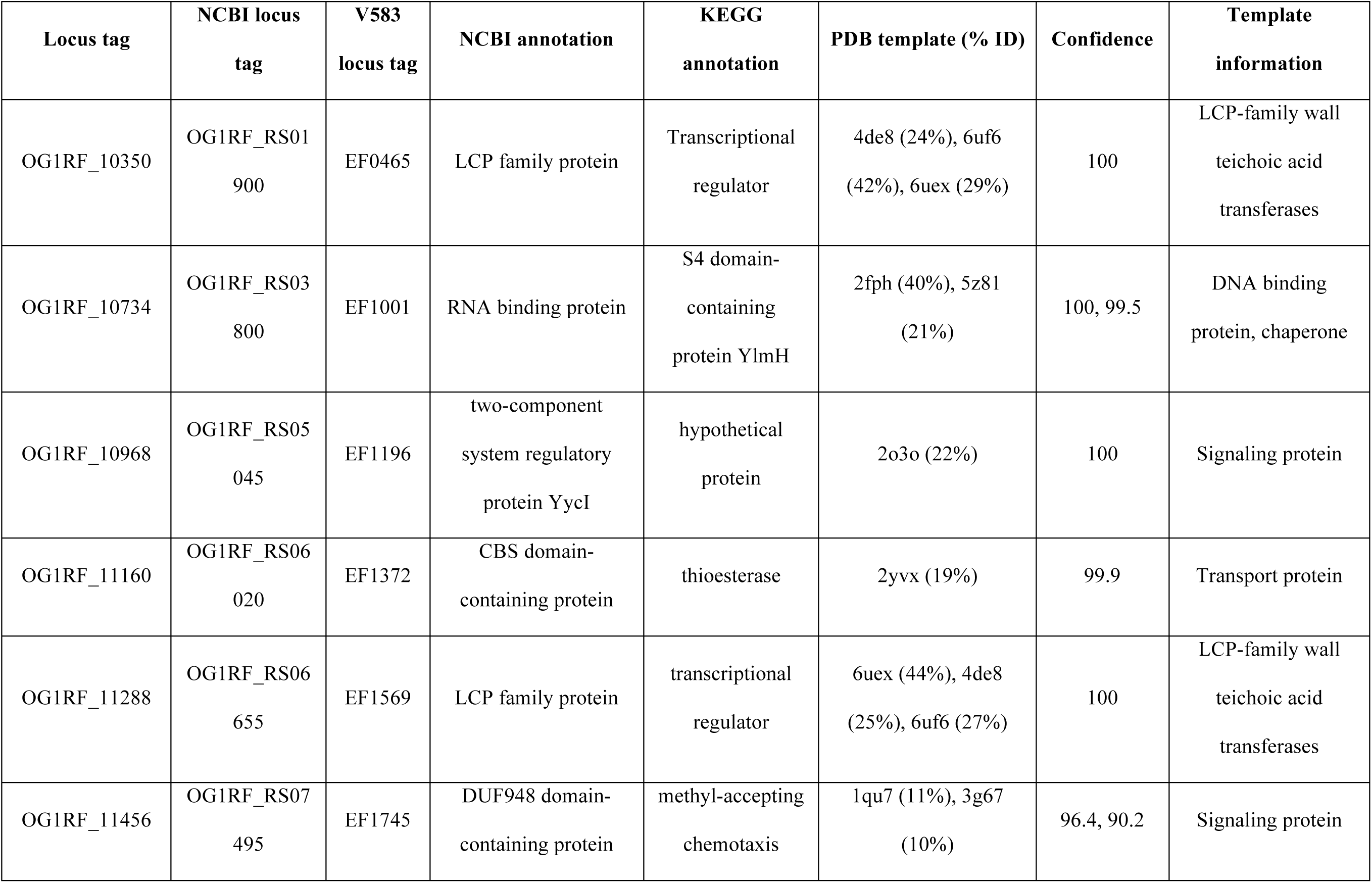
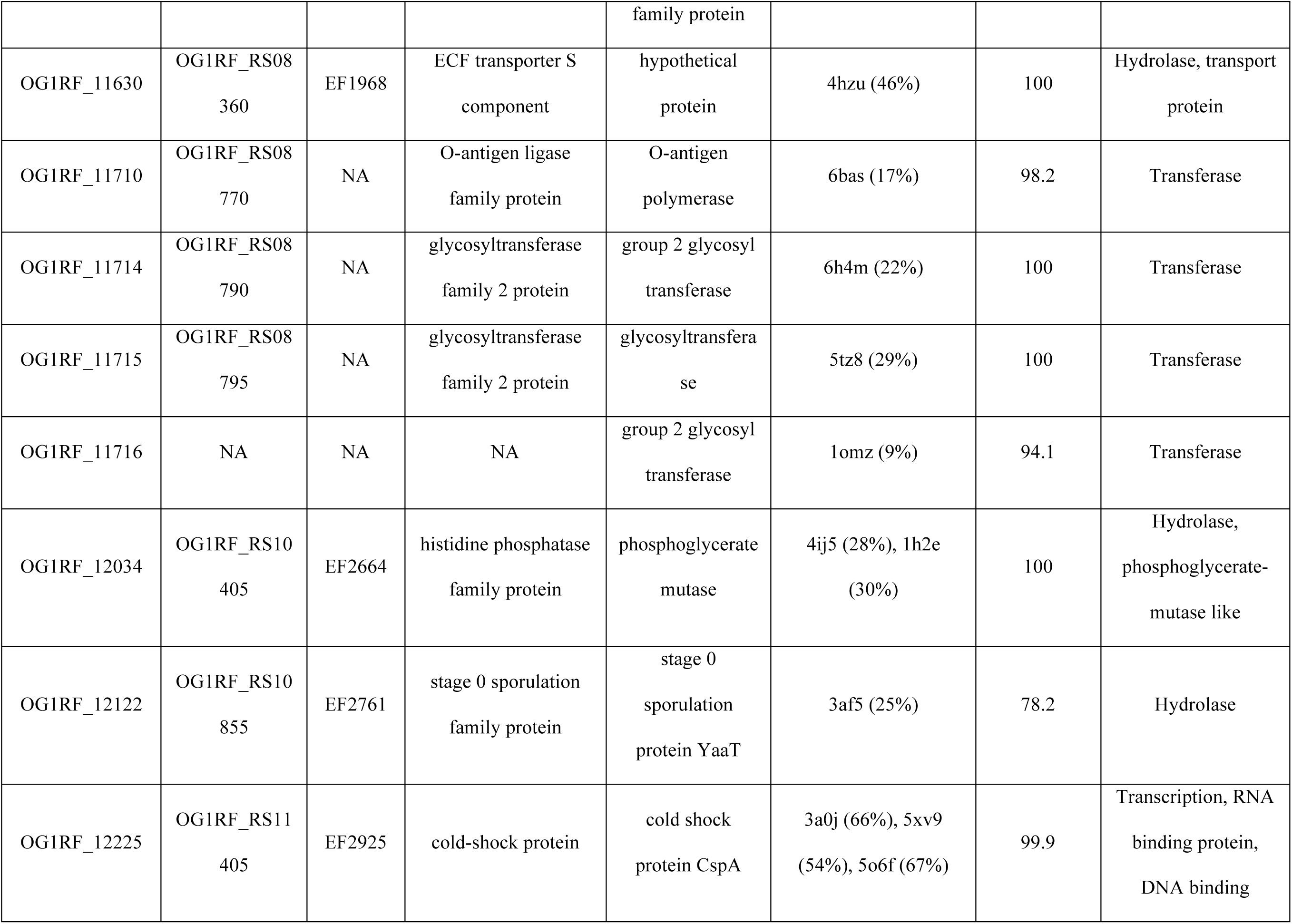
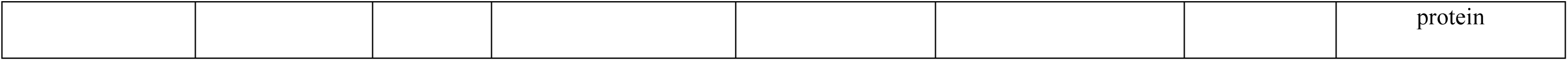
Structure and function predictions of poorly characterized biofilm determinants.

| **Locus tag** | **NCBI locus tag** | **V583 locus tag** | **NCBI annotation** | **KEGG annotation** | **PDB template (% ID)** | **Confidence** | **Template information** |
| --- | --- | --- | --- | --- | --- | --- | --- |
| OG1RF_10350 | OG1RF_RS01900 | EF0465 | LCP family protein | Transcriptional regulator | 4de8 (24%), 6uf6 (42%), 6uex (29%) | 100 | LCP-family wall teichoic acid transferases |
| OG1RF_10734 | OG1RF_RS03800 | EF1001 | RNA binding protein | S4 domain-containing protein YlmH | 2fph (40%), 5z81 (21%) | 100, 99.5 | DNA binding protein, chaperone |
| OG1RF_10968 | OG1RF_RS05045 | EF1196 | two-component system regulatory protein YycI | hypothetical protein | 2o3o (22%) | 100 | Signaling protein |
| OG1RF_11160 | OG1RF_RS06020 | EF1372 | CBS domain-containing protein | thioesterase | 2yvx (19%) | 99.9 | Transport protein |
| OG1RF_11288 | OG1RF_RS06655 | EF1569 | LCP family protein | transcriptional regulator | 6uex (44%), 4de8 (25%), 6uf6 (27%) | 100 | LCP-family wall teichoic acid transferases |
| OG1RF_11456 | OG1RF_RS07495 | EF1745 | DUF948 domain-containing protein | methyl-accepting chemotaxis family protein | 1qu7 (11%), 3g67 (10%) | 96.4, 90.2 | Signaling protein |
| OG1RF_11630 | OG1RF_RS08<br>360 | EF1968 | ECF transporter S<br>component | hypothetical<br>protein | 4hzu (46%) | 100 | Hydrolase, transport<br>protein |
| OG1RF_11710 | OG1RF_RS08<br>770 | NA | O-antigen ligase<br>family protein | O-antigen<br>polymerase | 6bas (17%) | 98.2 | Transferase |
| OG1RF_11714 | OG1RF_RS08<br>790 | NA | glycosyltransferase<br>family 2 protein | group 2 glycosyl<br>transferase | 6h4m (22%) | 100 | Transferase |
| OG1RF_11715 | OG1RF_RS08<br>795 | NA | glycosyltransferase<br>family 2 protein | glycosyltransfera<br>se | 5tz8 (29%) | 100 | Transferase |
| OG1RF_11716 | NA | NA | NA | group 2 glycosyl<br>transferase | 1omz (9%) | 94.1 | Transferase |
| OG1RF_12034 | OG1RF_RS10<br>405 | EF2664 | histidine phosphatase<br>family protein | phosphoglycerate<br>mutase | 4ij5 (28%), 1h2e<br>(30%) | 100 | Hydrolase,<br>phosphoglycerate-<br>mutase like |
| OG1RF_12122 | OG1RF_RS10<br>855 | EF2761 | stage 0 sporulation<br>family protein | stage 0<br>sporulation<br>protein YaaT | 3af5 (25%) | 78.2 | Hydrolase |
| OG1RF_12225 | OG1RF_RS11<br>405 | EF2925 | cold-shock protein | cold shock<br>protein CspA | 3a0j (66%), 5xv9<br>(54%), 5o6f (67%) | 99.9 | Transcription, RNA<br>binding protein,<br>DNA binding protein |

**Table S4.**
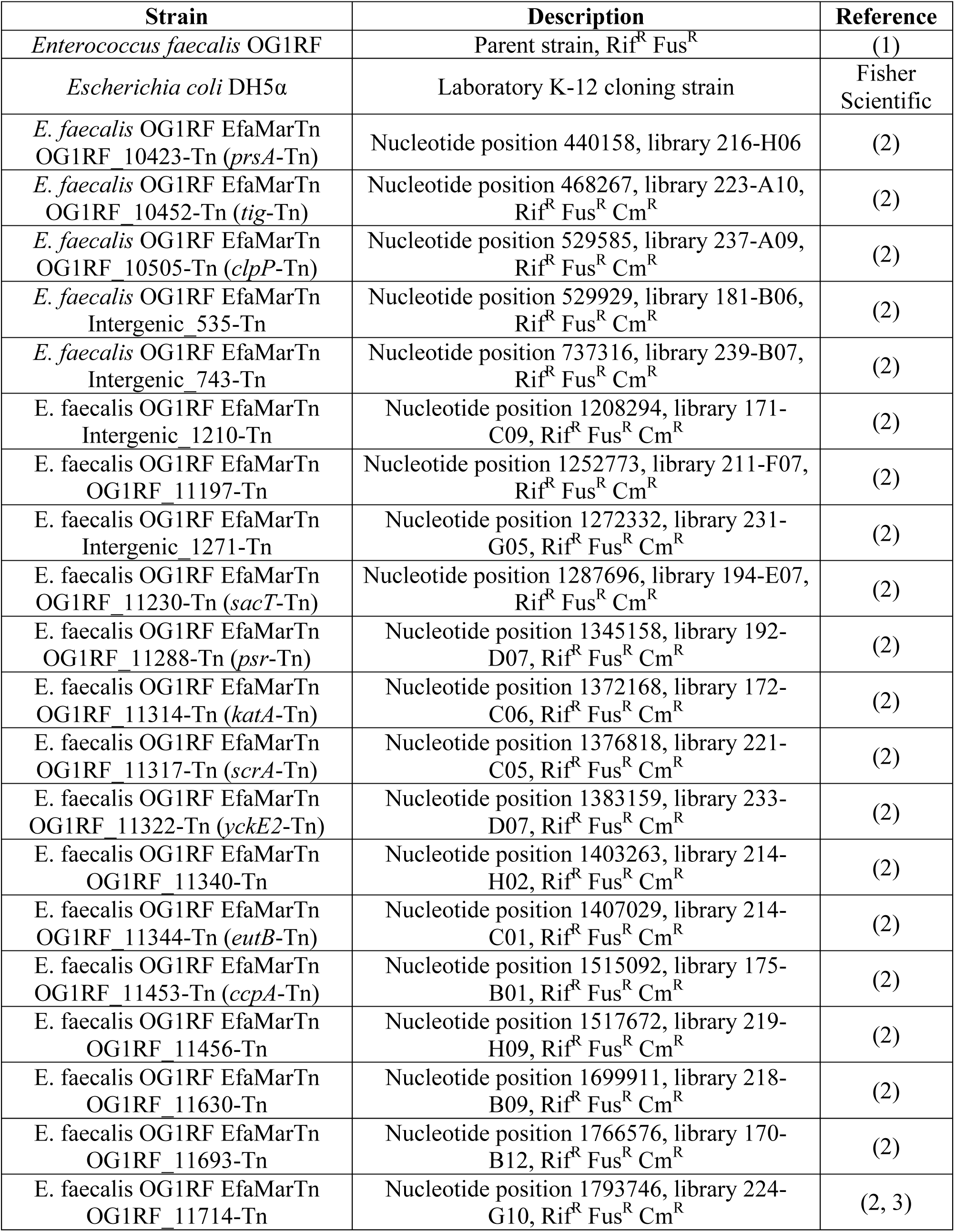
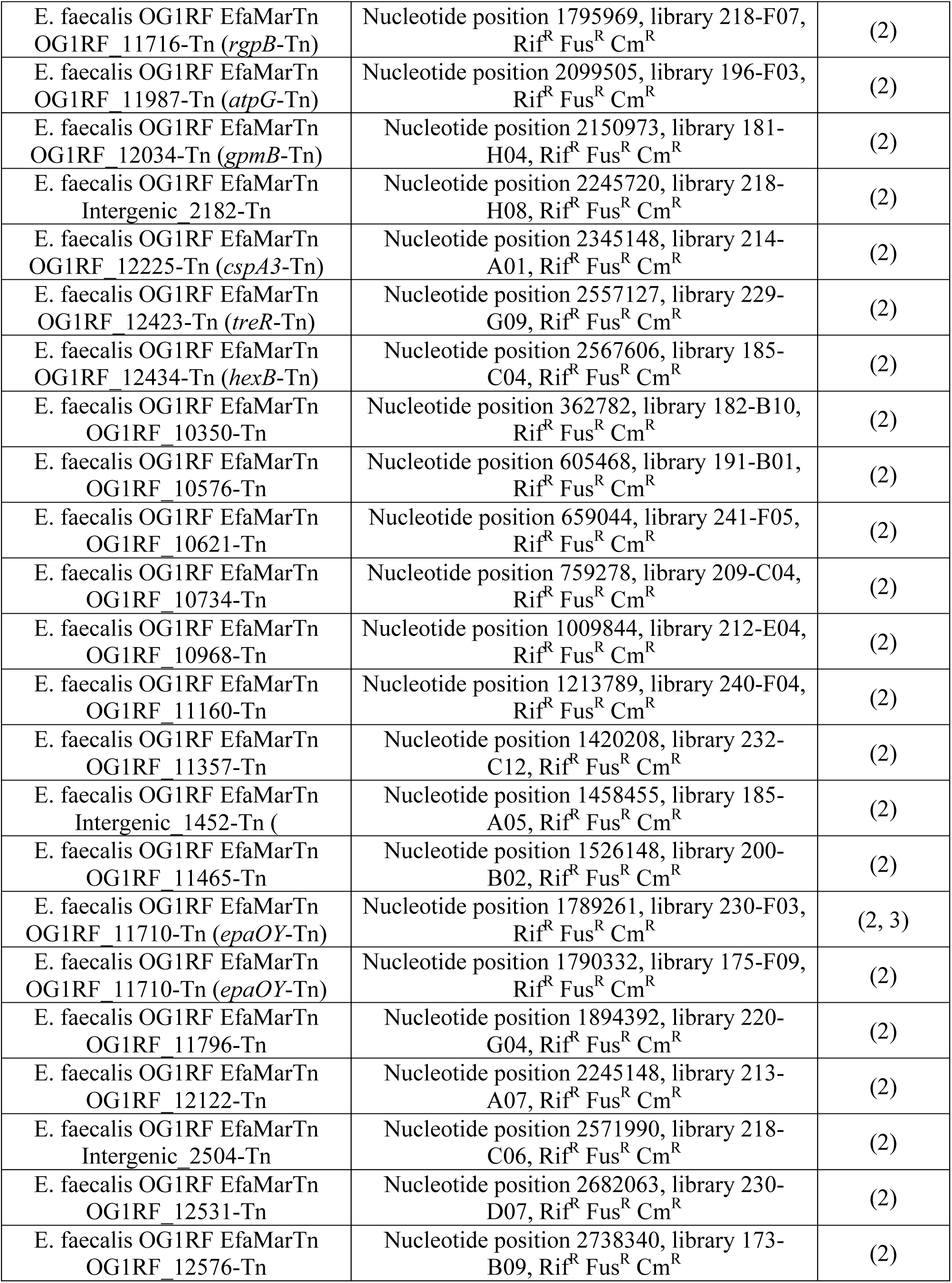
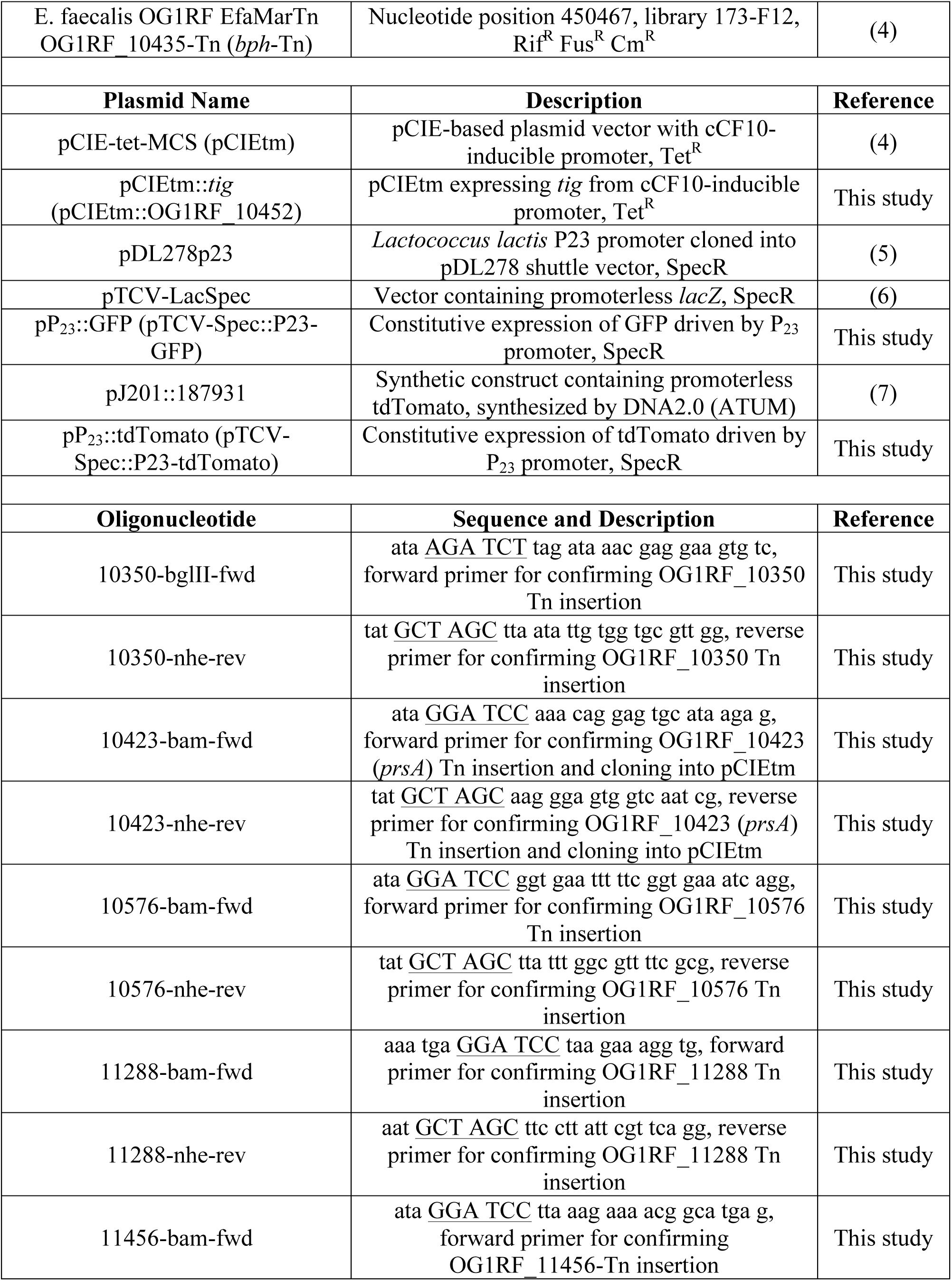
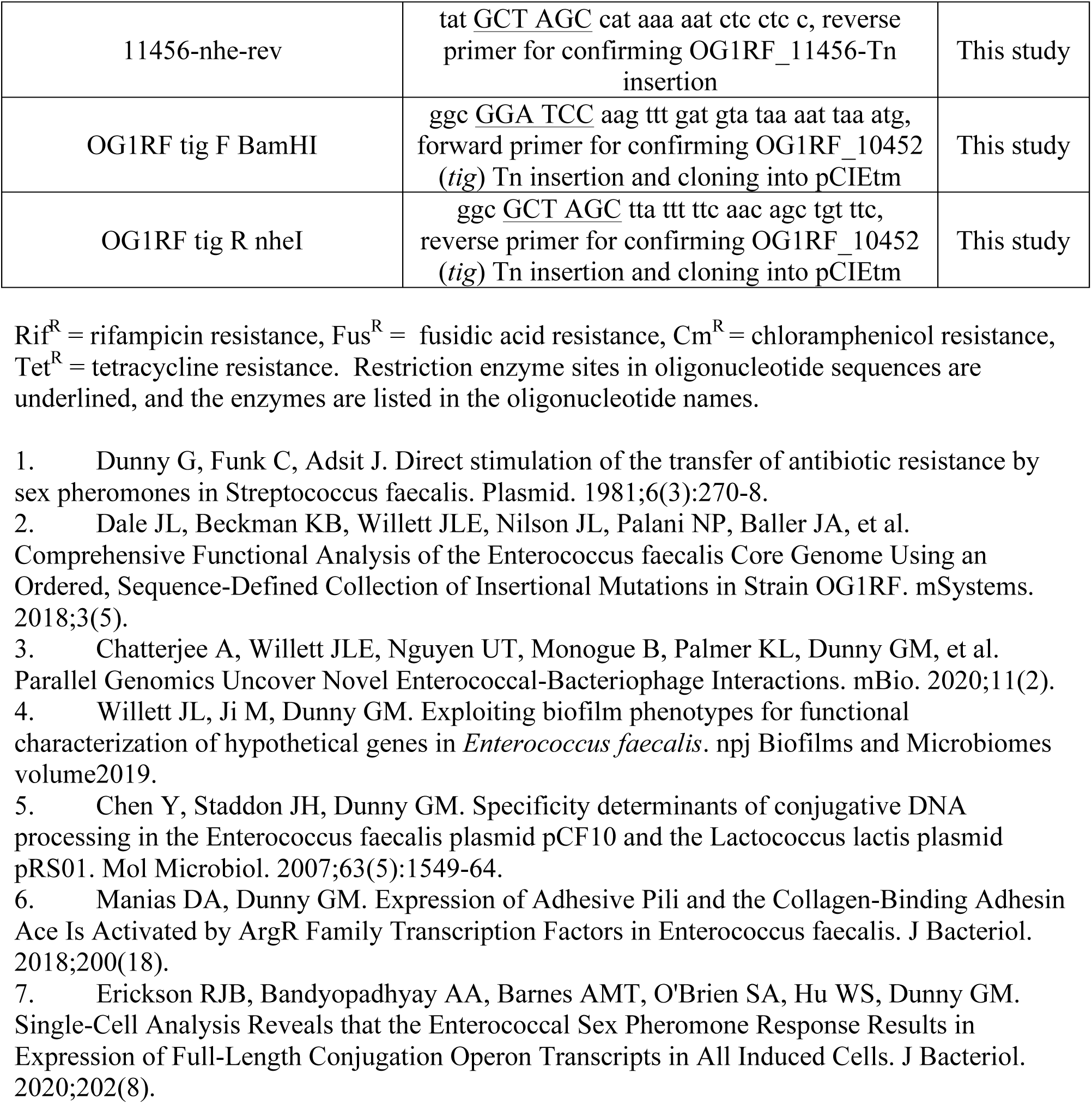
Strains, plasmids, and oligonucleotides used in this study.

## Notes

### Competing Interest Statement

The authors have declared no competing interest.

## References

1. Orrhage K, Nord CE. Factors controlling the bacterial colonization of the intestine in breastfed infants. Acta Paediatr Suppl. 1999;88(430):47–57.

2. Dubin K, Pamer EG. Enterococci and Their Interactions with the Intestinal Microbiome. Microbiol Spectr. 2014;5(6).

3. Schloissnig S, Arumugam M, Sunagawa S, Mitreva M, Tap J, Zhu A, et al. Genomic variation landscape of the human gut microbiome. Nature. 2013;493(7430):45–50.

4. Guiton PS, Hannan TJ, Ford B, Caparon MG, Hultgren SJ. Enterococcus faecalis overcomes foreign body-mediated inflammation to establish urinary tract infections. Infect Immun. 2013;81(1):329–39.

5. Ch’ng JH, Chong KKL, Lam LN, Wong JJ, Kline KA. Biofilm-associated infection by enterococci. Nat Rev Microbiol. 2019;17(2):82–94.

6. Madsen KT, Skov MN, Gill S, Kemp M. Virulence Factors Associated with Enterococcus faecalis Infective Endocarditis: A Mini Review. Open Microbiol J. 2017;11:1–11.

7. Wang QQ, Zhang CF, Chu CH, Zhu XF. Prevalence of Enterococcus faecalis in saliva and filled root canals of teeth associated with apical periodontitis. Int J Oral Sci. 2012;4(1):19–23.

8. Tornero E, Senneville E, Euba G, Petersdorf S, Rodriguez-Pardo D, Lakatos B, et al. Characteristics of prosthetic joint infections due to Enterococcus sp. and predictors of failure: a multi-national study. Clin Microbiol Infect. 2014;20(11):1219–24.

9. Keogh D, Tay WH, Ho YY, Dale JL, Chen S, Umashankar S, et al. Enterococcal Metabolite Cues Facilitate Interspecies Niche Modulation and Polymicrobial Infection. Cell Host Microbe. 2016;20(4):493–503.

10. Hollenbeck BL, Rice LB. Intrinsic and acquired resistance mechanisms in enterococcus. Virulence. 2012;3(5):421–33.

11. García-Solache M, Rice LB. The Enterococcus: a Model of Adaptability to Its Environment. Clin Microbiol Rev. 2019;32(2).

12. Gilmore MS, Lebreton F, van Schaik W. Genomic transition of enterococci from gut commensals to leading causes of multidrug-resistant hospital infection in the antibiotic era. Curr Opin Microbiol. 2013;16(1):10–6.

13. Lebreton F, Manson AL, Saavedra JT, Straub TJ, Earl AM, Gilmore MS. Tracing the Enterococci from Paleozoic Origins to the Hospital. Cell. 2017;169(5):849–61.e13.

14. Barnes AMT, Dale JL, Chen Y, Manias DA, Greenwood Quaintance KE, Karau MK, et al. Enterococcus faecalis readily colonizes the entire gastrointestinal tract and forms biofilms in a germ-free mouse model. Virulence. 2017;8(3):282–96.

15. Barnes AM, Ballering KS, Leibman RS, Wells CL, Dunny GM. Enterococcus faecalis produces abundant extracellular structures containing DNA in the absence of cell lysis during early biofilm formation. MBio. 2012;3(4):e00193–12.

16. Dale JL, Nilson JL, Barnes AMT, Dunny GM. Restructuring of Enterococcus faecalis biofilm architecture in response to antibiotic-induced stress. NPJ Biofilms Microbiomes. 2017;3:15.

17. Bjarnsholt T, Alhede M, Eickhardt-Sørensen SR, Moser C, Kühl M, Jensen P, et al. The in vivo biofilm. Trends Microbiol. 2013;21(9):466–74.

18. Merritt JH, Kadouri DE, O’Toole GA. Growing and analyzing static biofilms. Curr Protoc Microbiol. 2005;Chapter 1:Unit 1B.

19. Sankaran J, Karampatzakis A, Rice SA, Wohland T. Quantitative imaging and spectroscopic technologies for microbiology. FEMS Microbiol Lett. 2018;365(9).

20. Kristich CJ, Li YH, Cvitkovitch DG, Dunny GM. Esp-independent biofilm formation by Enterococcus faecalis. J Bacteriol. 2004;186(1):154–63.

21. Leuck AM, Johnson JR, Dunny GM. A widely used in vitro biofilm assay has questionable clinical significance for enterococcal endocarditis. PLoS One. 2014;9(9):e107282.

22. Dale JL, Beckman KB, Willett JLE, Nilson JL, Palani NP, Baller JA, et al. Comprehensive Functional Analysis of the Enterococcus faecalis Core Genome Using an Ordered, Sequence-Defined Collection of Insertional Mutations in Strain OG1R F. mSystems. 2018;3(5).

23. Willett JL, Ji M, Dunny GM. Exploiting biofilm phenotypes for functional characterization of hypothetical genes in Enterococcus faecalis. npj Biofilms and Microbiomes volume 2019.

24. Alhajjar N, Chatterjee A, Spencer BL, Burcham LR, Willett JLE, Dunny GM, et al. Genome-wide mutagenesis identifies factors involved in *Enterococcus faecalis* vaginal adherence and persistence. Infect Immun. 2020.

25. Chatterjee A, Willett JLE, Nguyen UT, Monogue B, Palmer KL, Dunny GM, et al. Parallel Genomics Uncover Novel Enterococcal-Bacteriophage Interactions. mBio. 2020;11(2).

26. Dale JL, Cagnazzo J, Phan CQ, Barnes AM, Dunny GM. Multiple roles for Enterococcus faecalis glycosyltransferases in biofilm-associated antibiotic resistance, cell envelope integrity, and conjugative transfer. Antimicrob Agents Chemother. 2015;59(7):4094–105.

27. Kristich CJ, Nguyen VT, Le T, Barnes AM, Grindle S, Dunny GM. Development and use of an efficient system for random mariner transposon mutagenesis to identify novel genetic determinants of biofilm formation in the core Enterococcus faecalis genome. Appl Environ Microbiol. 2008;74(11):3377–86.

28. Dunny GM, Clewell DB. Transmissible toxin (hemolysin) plasmid in Streptococcus faecalis and its mobilization of a noninfectious drug resistance plasmid. J Bacteriol. 1975;124(2):784–90.

29. Manias DA, Dunny GM. Expression of Adhesive Pili and the Collagen-Binding Adhesin Ace Is Activated by ArgR Family Transcription Factors in Enterococcus faecalis. J Bacteriol. 2018;200(18).

30. Eckert C, Lecerf M, Dubost L, Arthur M, Mesnage S. Functional analysis of AtlA, the major N-acetylglucosaminidase of Enterococcus faecalis. J Bacteriol. 2006;188(24):8513–9.

31. Qin X, Singh KV, Xu Y, Weinstock GM, Murray BE. Effect of disruption of a gene encoding an autolysin of Enterococcus faecalis OG1RF. Antimicrob Agents Chemother. 1998;42(11):2883–8.

32. Teng F, Singh KV, Bourgogne A, Zeng J, Murray BE. Further characterization of the epa gene cluster and Epa polysaccharides of Enterococcus faecalis. Infect Immun. 2009;77(9):3759–67.

33. Korir ML, Dale JL, Dunny GM. Role of e*paQ*, a Previously Uncharacterized Enterococcus faecalis Gene, in Biofilm Development and Antimicrobial Resistance. J Bacteriol. 2019;201(18).

34. Frank KL, Guiton PS, Barnes AM, Manias DA, Chuang-Smith ON, Kohler PL, et al. AhrC and Eep are biofilm infection-associated virulence factors in Enterococcus faecalis. Infect Immun. 2013;81(5):1696–708.

35. Hamoen LW, Meile JC, de Jong W, Noirot P, Errington J. SepF, a novel FtsZ-interacting protein required for a late step in cell division. Mol Microbiol. 2006;59(3):989–99.

36. Thomas VC, Hiromasa Y, Harms N, Thurlow L, Tomich J, Hancock LE. A fratricidal mechanism is responsible for eDNA release and contributes to biofilm development of Enterococcus faecalis. Mol Microbiol. 2009;72(4):1022–36.

37. Sillanpää J, Chang C, Singh KV, Montealegre MC, Nallapareddy SR, Harvey BR, et al. Contribution of individual Ebp Pilus subunits of Enterococcus faecalis OG1RF to pilus biogenesis, biofilm formation and urinary tract infection. PLoS One. 2013;8(7):e68813.

38. Nallapareddy SR, Singh KV, Sillanpää J, Garsin DA, Höök M, Erlandsen SL, et al. Endocarditis and biofilm-associated pili of Enterococcus faecalis. J Clin Invest. 2006;116(10):2799–807.

39. Crooke E, Wickner W. Trigger factor: a soluble protein that folds pro-OmpA into a membrane-assembly-competent form. Proc Natl Acad Sci U S A. 1987;84(15):5216–20.

40. Vorregaard M. Comstat2: a modern 3D image analysis environment for biofilms. Informatics and Mathematical Modelling: Technical University of Denmark.

41. Price MN, Wetmore KM, Waters RJ, Callaghan M, Ray J, Liu H, et al. Mutant phenotypes for thousands of bacterial genes of unknown function. Nature. 2018;557(7706):503–9.

42. Frank KL, Colomer-Winter C, Grindle SM, Lemos JA, Schlievert PM, Dunny GM. Transcriptome analysis of Enterococcus faecalis during mammalian infection shows cells undergo adaptation and exist in a stringent response state. PLoS One. 2014;9(12):e115839.

43. Abranches J, Tijerina P, Avilés-Reyes A, Gaca AO, Kajfasz JK, Lemos JA. The cell wall-targeting antibiotic stimulon of Enterococcus faecalis. PLoS One. 2014;8(6):e64875.

44. Kelley LA, Mezulis S, Yates CM, Wass MN, Sternberg MJ. The Phyre2 web portal for protein modeling, prediction and analysis. Nat Protoc. 2015;10(6):845–58.

45. Kawai Y, Marles-Wright J, Cleverley RM, Emmins R, Ishikawa S, Kuwano M, et al. A widespread family of bacterial cell wall assembly proteins. EMBO J. 2011;30(24):4931–41.

46. Eberhardt A, Hoyland CN, Vollmer D, Bisle S, Cleverley RM, Johnsborg O, et al. Attachment of capsular polysaccharide to the cell wall in Streptococcus pneumoniae. Microb Drug Resist. 2012;18(3):240–55.

47. Li FKK, Rosell FI, Gale RT, Simorre JP, Brown ED, Strynadka NCJ. Crystallographic analysis of. J Biol Chem. 2020;295(9):2629–39.

48. Kim KK, Yokota H, Kim SH. Four-helical-bundle structure of the cytoplasmic domain of a serine chemotaxis receptor. Nature. 1999;400(6746):787–92.

49. Pollard AM, Bilwes AM, Crane BR. The structure of a soluble chemoreceptor suggests a mechanism for propagating conformational signals. Biochemistry. 2009;48(9):1936–44.

50. Thurlow LR, Thomas VC, Narayanan S, Olson S, Fleming SD, Hancock LE. Gelatinase contributes to the pathogenesis of endocarditis caused by Enterococcus faecalis. Infect Immun. 2010;78(11):4936–43.

51. Hancock LE, Perego M. The Enterococcus faecalis fsr two-component system controls biofilm development through production of gelatinase. J Bacteriol. 2004;186(17):5629–39.

52. Reffuveille F, Connil N, Sanguinetti M, Posteraro B, Chevalier S, Auffray Y, et al. Involvement of peptidylprolyl cis/trans isomerases in Enterococcus faecalis virulence. Infect Immun. 2012;80(5):1728–35.

53. Guerardel Y, Sadovskaya I, Maes E, Furlan S, Chapot-Chartier MP, Mesnage S, et al. Complete Structure of the Enterococcal Polysaccharide Antigen (EPA) of Vancomycin-Resistant Enterococcus faecalis V583 Reveals that EPA Decorations Are Teichoic Acids Covalently Linked to a Rhamnopolysaccharide Backbone. mBio. 2020;11(2).

54. Rigottier-Gois L, Madec C, Navickas A, Matos RC, Akary-Lepage E, Mistou MY, et al. The surface rhamnopolysaccharide epa of Enterococcus faecalis is a key determinant of intestinal colonization. J Infect Dis. 2015;211(1):62–71.

55. Chen Y, Staddon JH, Dunny GM. Specificity determinants of conjugative DNA processing in the Enterococcus faecalis plasmid pCF10 and the Lactococcus lactis plasmid pRS01. Mol Microbiol. 2007;63(5):1549–64.

56. Breuer RJ, Bandyopadhyay A, O’Brien SA, Barnes AMT, Hunter RC, Hu WS, et al. Stochasticity in the enterococcal sex pheromone response revealed by quantitative analysis of transcription in single cells. PLoS Genet. 2017;13(7):e1006878.

57. Kristich CJ, Chandler JR, Dunny GM. Development of a host-genotype-independent counterselectable marker and a high-frequency conjugative delivery system and their use in genetic analysis of Enterococcus faecalis. Plasmid. 2007;57(2):131–44.

58. Schindelin J, Arganda-Carreras I, Frise E, Kaynig V, Longair M, Pietzsch T, et al. Fiji: an open-source platform for biological-image analysis. Nat Methods. 2012;9(7):676–82.

59. Heydorn A, Nielsen AT, Hentzer M, Sternberg C, Givskov M, Ersbøll BK, et al. Quantification of biofilm structures by the novel computer program COMSTAT. Microbiology. 2000;146 (Pt 10):2395–407.

60. Rossner M, Yamada KM. What’s in a picture? The temptation of image manipulation. J Cell Biol. 2004;166(1):11–5.

61. Krogh A, Larsson B, von Heijne G, Sonnhammer EL. Predicting transmembrane protein topology with a hidden Markov model: application to complete genomes. J Mol Biol. 2001;305(3):567–80.

62. The PyMOL Molecular Graphics System, Version 2.0. Schrödinger, LLC.

